# The phosphatase PTPRG controls FGFR1 activity and influences sensitivity to FGFR kinase inhibitors

**DOI:** 10.1101/120204

**Authors:** Michal Kostas, Ellen Margrethe Haugsten, Yan Zhen, Vigdis Sørensen, Patrycja Szybowska, Elisa Fiorito, Susanne Lorenz, Gustavo Antonio de Souza, Antoni Wiedlocha, Jørgen Wesche

## Abstract

FGFR1 represents an important target for precision medicine and a detailed molecular understanding of the target is important in order to increase the efficacy of FGFR inhibitors. We have here applied proximity labelling of FGFR1 in an osteosarcoma cell line to identify determinants of FGFR1 activity. Many known FGFR interactors were identified (e.g. FRS2, PLCγ, RSK2, SHC4, SRC), but the data also suggested novel determinants. A strong hit in our screen was the tyrosine phosphatase PTPRG. We show that PTPRG and FGFR1 interact and colocalize at the plasma membrane where PTPRG directly dephosphorylates activated FGFR1. We further show that osteosarcoma cell lines depleted for PTPRG display increased FGFR activity and are hypersensitive to stimulation by FGF1. In addition, PTPRG depletion elevated cell growth and negatively affected the efficacy of FGFR kinase inhibitors. Thus, PTPRG may have future clinical relevance by being a predictor of outcome after FGFR inhibitor treatment.

## Introduction

The fibroblast growth factor receptor (FGFR) family consists of four receptor tyrosine kinases (FGFR1-4), which are composed of an extracellular ligand binding part, a single transmembrane spanning stretch and an intracellular domain containing a tyrosine kinase (Turner and Grose, 2010, Haugsten et al., 2010). Upon ligand (FGF) binding, dimerization causes the receptors to auto-transphosporylate, leading to activation of downstream signalling cascades that regulate many key cellular responses such as proliferation, differentiation and cell migration. Importantly, aberrant FGF signalling is often involved in cancer development (Wesche et al., 2011, Turner and Grose, 2010). FGFR overexpression and activating mutations have recently been demonstrated to play an important role in several types of sarcoma (e.g. osteosarcoma, rhabdomyosarcoma (RMS) and soft tissue sarcoma) (Weekes et al., 2016, Guagnano et al., 2012, Taylor et al., 2009, Chudasama et al., 2016, Zhou et al., 2016). In addition, the FGFR-specific downstream signalling adaptor, the FGFR substrate 2 (FRS2), is overexpressed in liposarcoma and render these cells sensitive to FGFR inhibitors (Hanes et al., 2016, Zhang et al., 2013).

The incidence of sarcoma in adults is low (approx. 1% of all cancers), but more frequent in children and adolescents (approx. 10%) (Zhou et al., 2016). There is little commercial interest in these small and heterogeneous patient groups, and for the same reasons, they are difficult to investigate and it is challenging to develop better treatments. There are, however, several initiatives to develop drugs specific for FGFRs that possibly could also be used to treat sarcomas with aberrant FGFR signalling (Dieci et al., 2013). Most of these involve the development of specific small-molecular tyrosine kinase inhibitors and some have entered clinical trials for instance in patients with glioma, renal clear cell carcinoma, breast and lung cancer (ClinicalTrials.gov).

Unfortunately, in some cases such inhibitors fail even in the presence of the FGFR biomarker, for unknown reasons (Tabernero et al., 2015). There have also been reported effects of FGFR inhibitors in osteosarcoma cells without apparent FGFR aberrations, indicating that other mechanisms for FGFR vulnerability exists (Hanes et al., 2016). To increase the impact of FGFR inhibitors, it is crucial to understand in detail how their action on FGFR signalling and cell viability is determined.

As FGFR1 is overexpressed in 18.5% of osteosarcomas with poor response to chemotherapy and constitute a new and important therapeutic target for these patients (Fernanda Amary et al., 2014, Baroy et al., 2016), we wanted to better understand how FGFR signalling is regulated. We, therefore, took advantage of the BioID proximity biotinylation system to identify determinants of FGFR1 signalling in osteosarcoma cells (Roux et al., 2012). Using this approach, we discovered that the tyrosine phosphatase receptor type G (PTPRG) negatively regulates FGFR1 activation in osteosarcoma. Cells depleted for PTPRG exhibit increased activation of FGFR and are more sensitive in mitogenic responses to FGF stimulation. Thus, PTPRG seems to be important for controlling excessive FGFR signalling, which corresponds well with previous reports that implicate PTPRG as a tumour suppressor (LaForgia et al., 1991, Shu et al., 2010). Importantly, we found that PTPRG determines the sensitivity of cells to kinase inhibitors of FGFRs. We believe this may have clinical relevance as clinical cases with overexpressed FGFR1 combined with low expression of PTPRG have been reported.

## Results

### Proteomic BioID screen identifies determinants of FGFR1 activity

To identify proteins involved in the regulation of FGFR1 signalling in osteosarcoma cells, we performed a BioID screen by fusing a biotin ligase, BirA*, to the C-terminal tail of FGFR1. We recently validated and used this system to investigate signalling and trafficking of the related FGFR4 (Haugsten et al., 2016). When expressed in U2OS cells the biotin ligase biotinylates proteins in its proximity upon addition of biotin. The biotinylated proteins may be isolated by affinity to streptavidin and identified by quantitative LC-MS/MS. Since FGFR1-regulating proteins could be proximal to the receptor both in its inactive or its active state, we included samples of unstimulated and FGF1-stimulated U2OS cells stably expressing FGFR1 fused to BirA* (U2OS-R1-BirA*) (see S1 and S2 in Fig. 1A).

**Figure 1.**
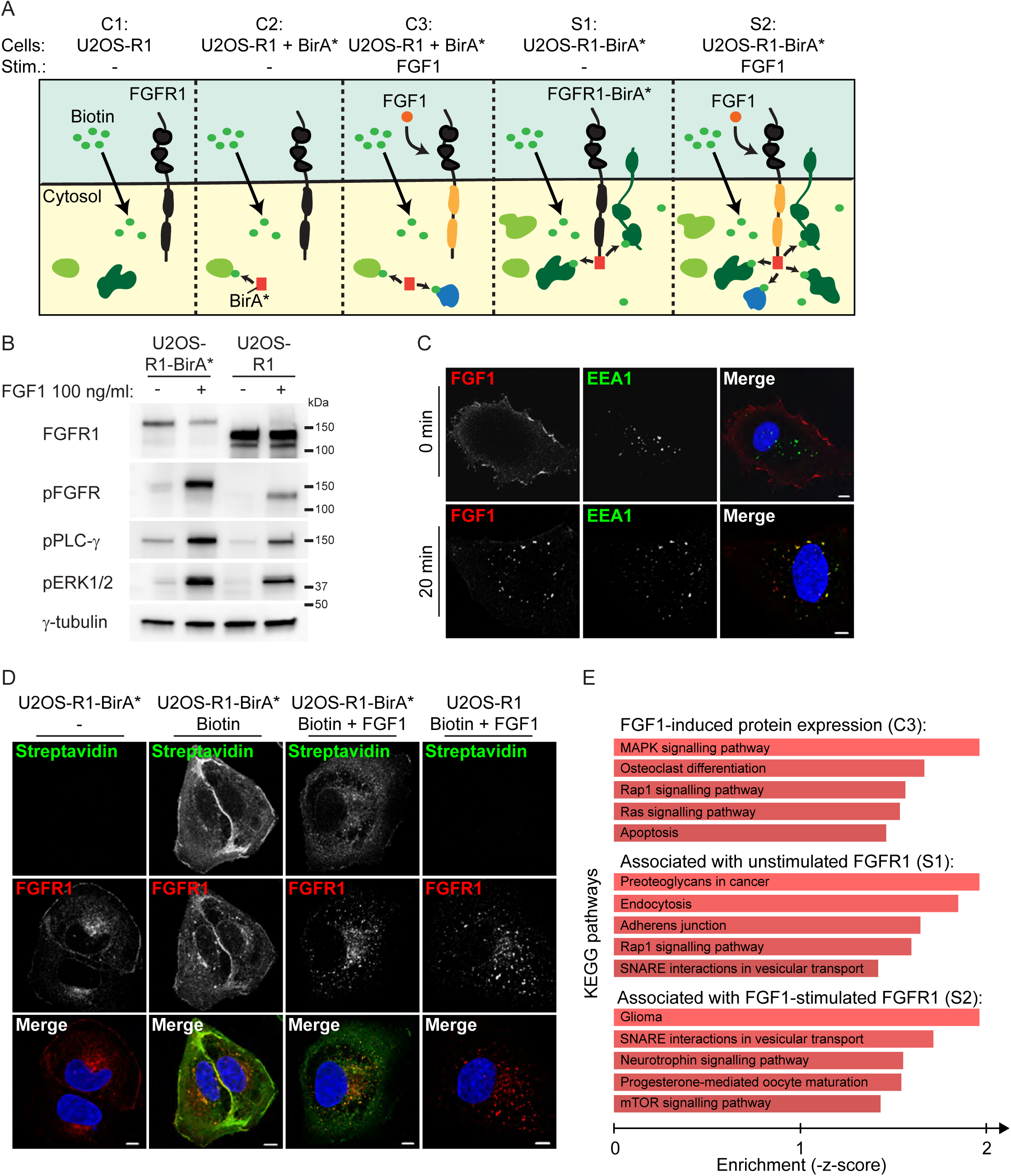
A BioID proteomic screen for determinants of FGFR1 activity in osteosarcoma cells. **(A)** A schematic presentation of the BioID experiment. Upon addition of biotin to cells, proteins in close proximity to the BirA* tag will be biotinylated. Biotinylated proteins are then isolated by Streptavidin pulldown and identified by quantitative LC MS/MS. The following five conditions are compared: C1 (Control 1): U2OS cells stably expressing wild-type FGFR1 (U2OS-R1), C2 (Control 2): U2OS cells stably coexpressing wild-type FGFR1 and BirA* (U2OS-R1 + BirA*), C3 (Control 3): U2OS cells stably coexpressing wild-type FGFR1 and BirA* stimulated with FGF1 (U2OS-R1 + BirA* + FGF1), S1 (Sample 1): U2OS cells stably expressing FGFR1 fused to BirA* (U2OS-R1-BirA*), S2 (Sample 2): U2OS cells stably expressing FGFR1 fused to BirA* (U2OS-R1-BirA*) and stimulated with FGF1. Biotin is added in all conditions. Addition of FGF1 induces activation of the receptor and its downstream signalling (indicated in yellow). Proteins identified in C1 and C2 represent the background while proteins identified in C3 represent proteins with induced expression upon FGF1 stimulation. Proteins identified in S1 represent proteins in proximity to unstimulated FGFR1 while proteins identified in S2 represents proteins in proximity to FGF1-activated FGFR1. Proteins in proximity to the receptor are indicated by dark green and proteins further away from the receptor are indicated in light green. Proteins with increased expression upon FGF1 stimulation are indicated in blue. **(B)** U2OS-R1 cells or U2OS-R1-BirA* cells were starved for 3 hours in serum free media before stimulation for 20 minutes with 100 ng/ml FGF1 in the presence of heparin (20 U/ml). Cells were then lysed and the cellular material was analysed by SDS-PAGE and western blotting using the indicated antibodies. A p in front of the name of the antibody indicates that it recognizes the phosphorylated form of the protein. Note that the total FGFR1 antibody recognizes the tagged version of FGFR1 (FGFR1-BirA*) less efficient than the wild-type receptor. A representative western blot is shown. **(C)** U2OS-R1-BirA* cells were allowed to bind DL550-FGF1 at 4°C in the presence of heparin and then washed (to remove excess DL550-FGF1) and either fixed directly (0 min) or incubated for 20 min at 37°C before fixation (20 min). Fixed cells were stained with anti-EEA1 antibody and Hoechst and examined by confocal microscopy. Representative cells are shown. Scale bar 5 μm. **(D)** U2OS-R1-BirA* and U2OS-R1 cells were either left untreated or treated with 50 mM biotin and/or 100 ng/ml FGF1 in the presence of 10 U/ml heparin as indicated for 24 hours. The cells were then fixed and stained with anti FGFR1 antibody, Alexa 488 streptavidin and Hoechst. Merged images are shown in the bottom panel. Representative cells are shown. Scale bar 5 μm. **(E)** KEGG pathways analyses were applied to the three datasets using Enrichr (http://amp.pharm.mssm.edu/Enrichr/)(Kuleshov et al., 2016).

Control samples included U2OS cells stably expressing FGFR1 wild-type (U2OS-R1, C1 in Fig. 1A) and U2OS cells stably co-expressing FGFR1 wild-type and control, non-fused BirA* (U2OS-R1 + BirA*, C2 in Fig. 1A). Proteins identified in these two conditions were considered as background. As FGFR signalling induces increased expression of certain proteins, increased expression of background proteins in FGF stimulated conditions could erroneously be considered as positive hits. We therefore also included samples of U2OS cells stably co-expressing FGFR1 wild-type and BirA* (U2OS-R1 + BirA*) and stimulated with FGF1 (C3 in Fig. 1A). Such proteins were therefore subtracted from the final list in FGF stimulated conditions.

We first investigated whether the fusion of BirA* to FGFR1 could interfere with its functionality (Fig. 1B). After 20 minutes stimulation with FGF1, we detected comparable phosphorylation of the receptors itself, as well as known downstream signalling molecules, such as PLC-γ and ERK1/2. Because, the total FGFR1 antibody used for these western blots recognizes the wild-type receptor better than the tagged version, the staining underestimated the level of FGFR1-BirA* in these cells. Next, we tested whether the FGFR1-BirA* fusion protein is able to bind its ligand FGF1 at the cell surface and undergo endocytosis (Fig. 1C). Cells were kept on ice to facilitate binding of fluorophore-labelled FGF1 (DL550-FGF1) and heated to 37°C to allow internalization. Then, the cells were stained and examined by confocal microscopy. DL550-FGF1 was clearly detected at the cell surface in cells kept on ice and next, after heating to 37°C for 20 minutes, it was detected in intracellular structures colocalizing with EEA1. The results demonstrate that FGFR1-BirA* is able to bind FGF1 and internalize into early endosomes similarly to wild-type receptor (Haugsten et al., 2005).

Next, we tested the biotinylation efficiency of the fusion protein (Fig. 1D and Fig. 1 supplement 1). In the absence of biotin, little biotinylated proteins were detected in cells expressing FGFR1-BirA*. In the presence of biotin, a smear of bands representing biotinylated proteins was detected on western blotting (Fig. supplement 1) and a strong streptavidin staining was detected in cells by confocal microscopy (Fig. 1D). The streptavidin staining was stronger at the cell periphery close to the plasma membrane. When cells were treated with FGF1, the streptavidin staining was more dispersed in the cytoplasm, probably reflecting the transport of the receptor from the plasma membrane to endosomes upon FGF1 stimulation. Biotinylated proteins were barely visible in cells expressing FGFR1 wild-type and treated with biotin. Taken together these data indicate that the FGFR1-BirA* fusion protein is functional and active and efficiently biotinylates proteins in its proximity in the presence of biotin.

In the U2OS cells stably co-expressing non-fused BirA* and FGFR1 wild-type, a smear of bands representing biotinylated proteins were detected on western blotting (Fig. supplement 1) indicating that the biotin ligase is active in these cells. These cells express higher amounts of control BirA* than those expressing the FGFR1-BirA* fusion protein, which was an advantage to eliminate false positive hits in the screen.

Next, we performed the BioID proximity labelling and analysed affinity-purified biotinylated proteins by LC-MS/MS (Fig. 1A). Since control BirA* is expressed uniformly in the cytosol and the nucleus, it biotinylates proteins in general and by comparing the control conditions with and without FGF1 stimulation, we could obtain an overview of which proteins are induced by FGF signalling in these cells. Proteins significantly enriched more than 10X (p<0.05) in C3 compared to C1 and C2, were considered as proteins with increased expression upon FGF1 signalling (Table S1). Top hits among these were proteins known to be induced by FGF1 signalling, such as the transcription factors JUNB, FOSL1, FOSL2 and JUND (See STRING interaction network, Fig. 1 Supplement 2) (Szklarczyk et al., 2015). Analysing the data using Enrichr and KEGG Pathways applications (Kuleshov et al., 2016), we find enrichment of signalling pathways and, interestingly, induction of Osteoclast differentiation (Fig. 1E), which reflect the cell context of our analysis (osteoesarcoma).

Proteins significantly enriched more than 10X (p<0.05) in S1 compared to C1 and C2 (Fig 1A) were considered as proteins in proximity of non-stimulated FGFR1 (Table S1), while proteins significantly enriched more than 10X (p<0.05) in S2 compared to C1, C2 and C3 and, in addition enriched compared to S1, were considered as proteins in the proximity of the active receptor (Table S1). Among the proteins enriched in samples of the activated receptor, we identified several well-known FGFR downstream signalling proteins (PLCγ, RSK2 and SHC4) (Mohammadi et al., 1991, Nadratowska-Wesolowska et al., 2014), thereby validating our approach. FRS2 is constitutively bound to FGFR1 (Kouhara et al., 1997) and was accordingly found to be in the proximity of both unstimulated and stimulated receptors. Also SRC, previously found to be important for FGFR signalling and trafficking (Sandilands et al., 2007), was found associated with both unstimulated and activated receptors (See STRING interaction networks Fig. 1 supplements 3 and 4).

In the case of unstimulated receptors, the hits were enriched for plasma membrane functions (proteoglycans and adherens junctions) and membrane transport (endocytosis and SNARE interactions) (Fig. 1E), reflecting the known plasma membrane localization and importance of trafficking of the receptors. Stimulated receptors also showed enrichment for membrane transport, but also several signalling pathways (Fig. 1E), as expected.

A very strong hit in our screen was the tyrosine phosphatases PTPRG (Sorio et al., 1995). Because PTPRG has previously been suggested to be a tumour suppressor (LaForgia et al., 1991), we chose to focus our attention to the possible regulatory role of PTPRG on FGFR1. Interestingly, PTPRG was also found associated with both unstimulated and activated receptor.

### PTPRG interacts and co-localizes with FGFR1 at the plasma membrane

To validate the interaction between PTPRG and FGFR1 that was suggested by the BioID screen, we attempted co-immunoprecipitation of the two proteins. FGFR1-BirA* is fused to an HA-tag and U2OS-FGFR1-BirA* cells were transfected with MYC-FLAG-tagged PTPRG, lysed, and then immunoprecipitated with anti-HA antibodies and immunoblotted with anti-FLAG antibodies. The results demonstrated that PTPRG can efficiently be co-immunoprecipitated with FGFR1 indicating a physical interaction between PTPRG and FGFR1 (Fig. 2A).

**Figure 2.**
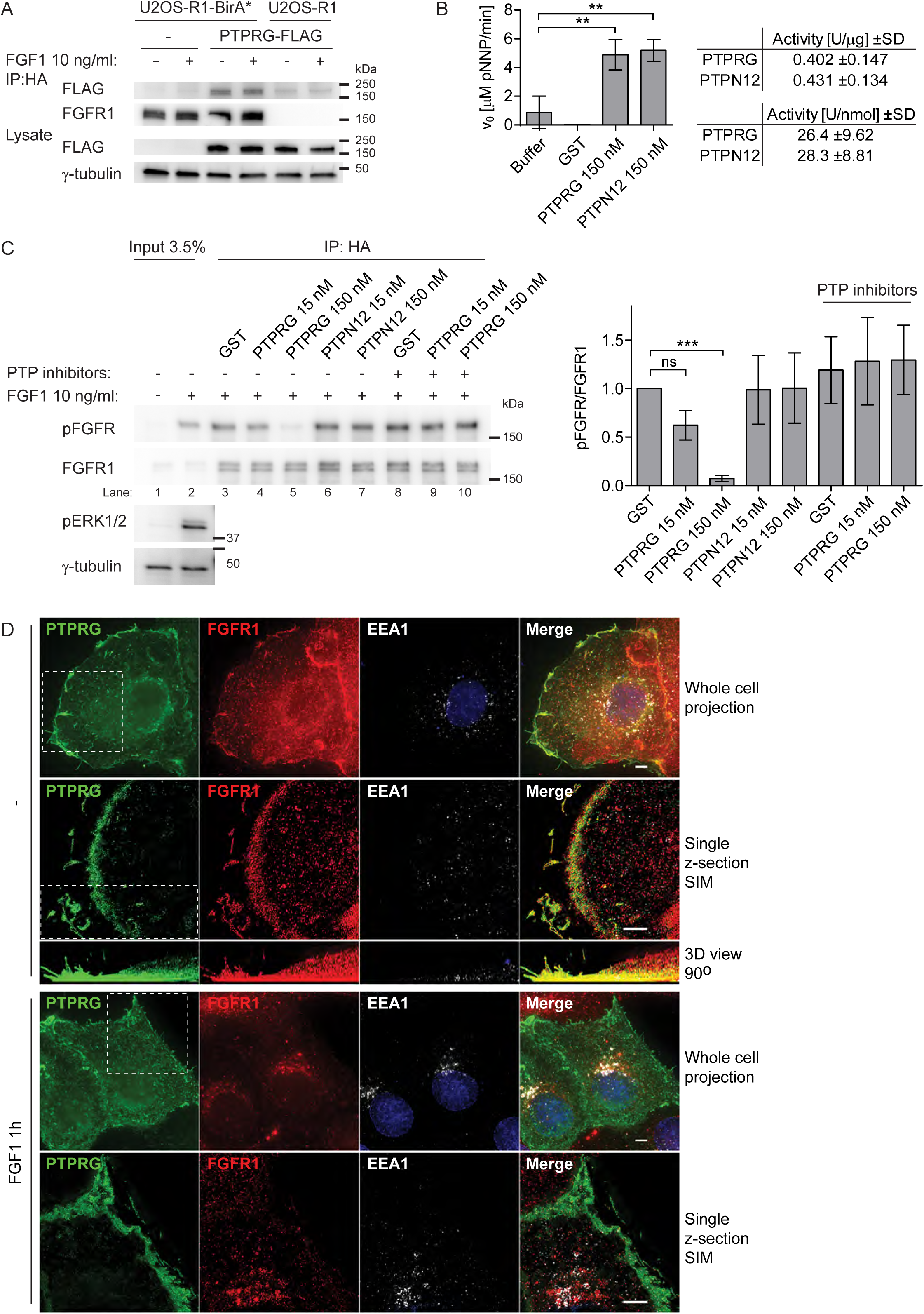
PTPRG binds and dephosphorylates FGFR1. **(A)** U2OS-R1-BirA* and U2OS-R1 cells were transfected with PTPRG-MYC-FLAG plasmid for 24 hours. U2OS-R1-BirA* cells not transfected with PTPRG were included as a control. Cells were then starved for 2 hours and left untreated or treated with 10 ng/ml FGF1 in the presence of 20 U/ml heparin for 15 minutes. After that, the cells were lysed and the lysates were subjected to immunoprecipitation using anti-HA magnetic beads followed by SDS-PAGE and western blotting with indicated antibodies. R1-BirA* is fused to an HA-tag in the C-terminal end. A representative western blot is shown. **(B)** Phosphatase activity of recombinant GST-PTPRG (catalytic domain) and GST-PTPN12 (catalytic domain) was estimated using pNPP assay. 2-fold molar excess of GST was used as a control. The initial rate of pNPP hydrolysis was measured colorimetrically (Abs. at 405 nm) during the first 10 min of reaction. The graph and table represent the mean ± SD of three independent experiments. **(C)** U2OS-R1-BirA* cells were serum-starved for 2 hours and then treated with 10 ng/ml FGF1 in the presence of 10 U/ml heparin for 15 minutes, lysed and the lysate was subjected to immunoprecipitation using anti-HA-tag antibodies. R1-BirA* is fused to an HA-tag in the C-terminal end. The beads containing immunoprecipitated FGFR1-BirA* were washed with lysis buffer without phosphatase inhibitor and subjected to on-beads dephosphorylation using indicated phosphatases or GST for 45 min, in the presence or absence of phosphatase inhibitors. After the incubation with phosphatases the immunoprecipiteted receptors were released from the beads and analysed by SDS-PAGE followed by western blotting using anti-pFGFR (Y653/Y654) antibodies. Western blots were quantified and bands corresponding to phosphorylated FGFR1 (pFGFR) were normalized to total FGFR1 immunoprecipitated and presented as fraction of GST without phosphatase inhibitors. The graph represents the mean ± SD of three independent experiments. The data were analysed using one-way RM ANOVA followed by Tukey *post hoc* test. ***p ≤ 0.001, ns - not-significant. **(D)** U2OS-R1 cells were transfected with PTPRG-MYC-FLAG, starved for 2 hours and either left untreated or treated with 200 ng/ml FGF1 and 10 U/ml heparin for 1 hour. The cells were fixed and stained with anti-FLAG, anti-FGFR1, anti-EEA1 antibodies and fluorophore (AF488, AF568, or AF647) labeled secondary antibodies and Hoechst. The cells were imaged in conventional wide-field mode and by SIM. Shown are Maximum Intensity Projections of whole cells (all z-sections) for deconvolved wide-field images, and a single selected optical section for SIM images, while all SIM z-sections were used for the 3D volume view, which was rotated 90° (side-view). Stippled lined squares indicate a region of the cell that is shown in a different mode in the panel below. Representative cells are shown. Scale bars 4 μm.

Since PTPRG protein contains an active phosphatase domain, we considered FGFR1 as a potential substrate for PTPRG. To test this, we performed an *in vitro* phosphatase assay using activated FGFR1, which was immunoprecipitated from FGF1-treated U2OS-R1-BirA* cell lysates, and a recombinant PTPRG phosphatase domain in fusion with GST. As a control in the experiment, we also used the recombinant phosphatase domain of PTPN12, which has been shown to dephosphorylate other receptor tyrosine kinases (Sun et al., 2011), and exhibited phosphatase activity towards a non-specific substrate (p-nitrophenyl phosphate, pNPP) comparable to PTPRG (Fig. 2B). After 45 min incubation with 150 nM PTPRG we observed a significant decrease in the level of FGFR1 phosphorylation at Y653/Y654 residues, as detected by western blotting, compared to incubation with 2-fold molar excess of GST (Fig. 2C, lane 5 compared to lane 3). Moreover, the dephosphorylation effect was less pronounced (and statistically insignificant) when we used 10-times lower concentration of the enzyme (Fig. 2C, lane 4), showing that the reduction in phospho-FGFR1 level is dependent on PTPRG concentration. We observed no changes in the phosphorylation of Y653/Y654 residues in FGFR1 using either 150 nM or 15 nM of PTPN12 (Fig. 2C, lane 6-7), suggesting a substrate specificity in the dephosphorylation of FGFR1 by PTPRG. Moreover, the dephosphorylation effect was not visible in the presence of a tyrosine-phosphatase inhibitor cocktail (Fig. 2C, lane 9-10), confirming that the dephosphorylation directly relies on PTPRG enzymatic activity.

To gain insight into where in the cells PTPRG and FGFR1 interact, we used wide-field and structured illumination microscopy to investigate their co-localization. In U2OS-R1 cells expressing PTPRG, FGFR1 and PTPRG co-localized mainly at the plasma membrane in non-stimulated cells (Fig. 2D). Interestingly, at resting conditions FGFR1 and PTPRG strongly co-localized in protrusions of the cells, resembling filopodia and lamellopodia. When cells had been stimulated with FGF1 for one hour, FGFR1 was detected mainly in intracellular vesicular structures, including EEA1 positive endosomes. PTPRG, however, was predominantly observed at the cell surface also after stimulation, suggesting that PTPRG might mainly act on FGFR1 at the plasma membrane (Fig. 2D).

### Regulation of FGFR1 autophosphorylation by PTPRG revealed by TIRF

PTPRG is a large, transmembrane tyrosine phosphatase with an extracellular part containing a carbonic anhydrase-like (CAH) domain and an intracellular part consisting of one active and one inactive phosphatase domain (Fig. 3A) (Barnea et al., 1993). Mutating the aspartate (D) at positon 1028 to alanine (A) inactivates the phosphatase activity of PTPRG (Zhang et al., 2012).

**Figure 3.**
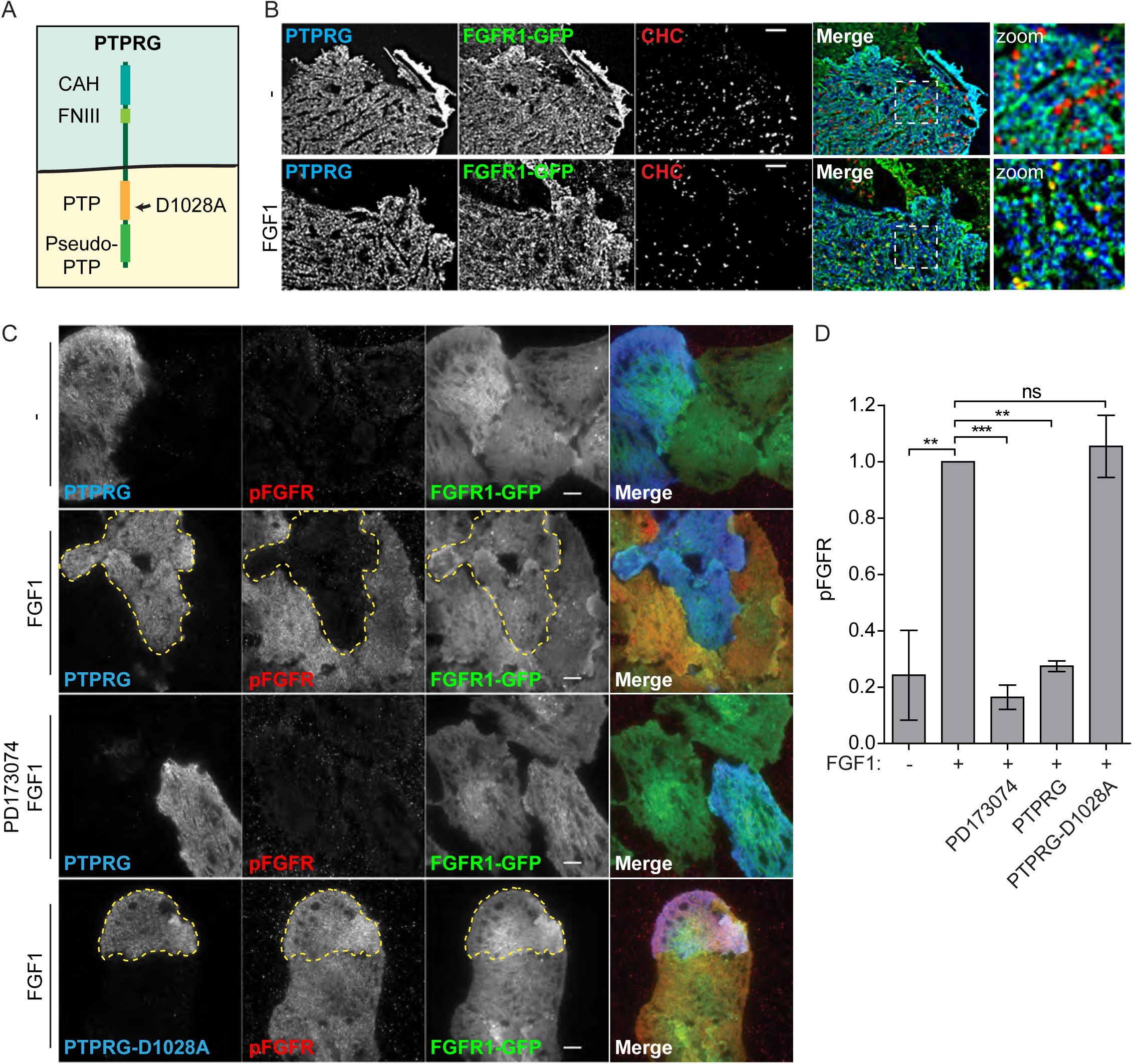
PTPRG counters FGFR1 autophosphorylation. **(A)** Schematic presentation of PTPRG. PTPRG is a transmembrane protein with an extracellular carbonic anhydrase-like domain (CAH) and a fibronectin type III-like domain (FNIII). The intracellular part contains two protein tyrosine phosphatase domains (PTP) of which only one is active (indicated in orange). The other is called a pseudo-PTP. Mutation of aspartic acid 1028 to alanine inactivates the phosphatase activity. **(B)** U2OS-R1-GFP cells transfected with PTPRG-myc-FLAG (for 20 hours), serum starved for 2 hours, and unstimulated (-) or stimulated with FGF1 for 15 min (FGF1), were fixed and stained with anti-myc and anti-Clathrin heavy chain, and imaged by TIRF. Merged images are overlays of PTPRG in blue, FGFR-GFP in green, and Clathrin in red. Blue and green overlay appears cyan. Green and red overlay appears yellow. Images were deconvolved and representative cells are shown. Scale bar 4 μm.**(C)** U2OS-R1-GFP cells were transfected with MYC-FLAG-tagged PTPRG or PTPRG-D1028A (for 20 hours), starved for 2 hours, and stimulated (or not) with FGF1 in the presence of heparin for 10 minutes (in one case in the presence of FGFR1 tyrosine kinase inhibitor PD173074) and then fixed and stained with anti-FLAG, anti-pFGFR1 (Y653/Y654), and fluorophore labeled secondary antibodies. The cells were imaged by TIRF and representative cells are shown. Merged images are overlays of PTPRG in blue, pFGFR1 in red, and FGFR1-GFP in green. Stippled lines indicate cells transfected with PTPRG or PTPRG-D1028A. Scale bars 8 μm. **(D)** The signal intensities for pFGFR1 in PTPRG-transfected or -untransfected cells were measured for 15-30 cells for each condition in three independent experiments and is presented as the mean values ± SD where values had been normalized to the signal intensity of untransfected cells stimulated with FGF1. The data were analysed using one-way RM ANOVA followed by Tukey *post hoc* test. ***p ≤ 0.001, **p ≤ 0.01, ns - not-significant.

Imaging by total internal reflection microscopy (TIRF), an imaging technique that reveals with high selectivity and clarity structures on, or close to, the cell surface, confirmed the localization of PTPRG at the plasma membrane. We observed a high degree of colocalization with FGFR1, but not with Clathrin Heavy Chain marking clathrin coated pits, which are entry sites for endocytosis (Fig. 3B, upper panel). Upon FGF1 stimulation, a partial shift of FGFR1 into clathrin coated pits was observed. This was not observed for PTPRG, suggesting that PTPRG is not co-endocytosed with FGFR1 (Fig. 3B lower panel).

Our biochemical analyses suggested that PTPRG acts directly to dephosphorylate FGFR1 and the plasma membrane localization of PTPRG suggests that it might do so mainly at the plasma membrane. To test this in cells, we used TIRF to monitor autophosphorylated FGFR1 at the plasma membrane. In this experiment, U2OS cells stably expressing FGFR1-GFP (U2OS-R1-GFP) were briefly stimulated with FGF1 before fixation and the activated FGFR1 was detected with anti-FGFR phospho-Tyr653/654 specific antibodies. The stimulation with FGF1 was sufficient to induce a robust activation of FGFR1 (pFGFR1) at the plasma membrane, which was fully inhibited by the FGFR kinase inhibitor PD173074, demonstrating that the observed immunofluorescent signal was specific (Fig. 3C). The cells were also transiently transfected with PTPRG, which had a dramatic effect, almost completely inhibiting the activity of FGFR1 at the plasma membrane. Overexpression of the inactive mutant PTPRG-D1028A however, had no effect on the FGFR1 activity (Fig. 3C). Quantification of the pFGFR1 levels detected by TIRF imaging, showed that overexpression of PTPRG reduced pFGFR1 levels by at least 70% (Fig. 3D). These experiments demonstrate that the enzymatic phosphatase activity of PTPRG counter the autophosphorylation of FGFR1.

We then depleted PTPRG in U2OS-R1 cells using siRNA oligonucleotides, which were shown to efficiently deplete PTPRG (Fig. 4A and B). We also constructed PTPRG rescue mutants for siRNA oligo #1 that were resistant to siRNA depletion (PTPRG siRes #1 and PTPRG-D1028A siRes #1), (Fig. 4B). When PTPRG was knocked down, we observed a substantial increase in the levels of phosphorylated FGFR1 upon FGF1 stimulation (Fig. 4C). Moreover, using TIRF microscopy we could observe a similar effect at the plasma membrane (Fig. 4D and E). Depletion of PTPRG led to a strong increase in FGF-induced phosphorylation of FGFR1. This demonstrates that endogenous levels of PTPRG negatively regulate FGFR1 autophosphorylation. This effect could be totally reversed by transfecting the cells with the siRNA-resistant version of PTPRG, while the siRNA resistant version of the inactive mutant PTPRG-D1028A was not able to reverse the effect (Fig. 4D and E). Quantification of the pFGFR1 levels detected by TIRF imaging, showed that PTPRG knockdown increased pFGFR1 levels at least twofold (Fig. 4E), demonstrating that PTPRG is a highly efficient phosphatase for activated FGFR1.

**Figure 4.**
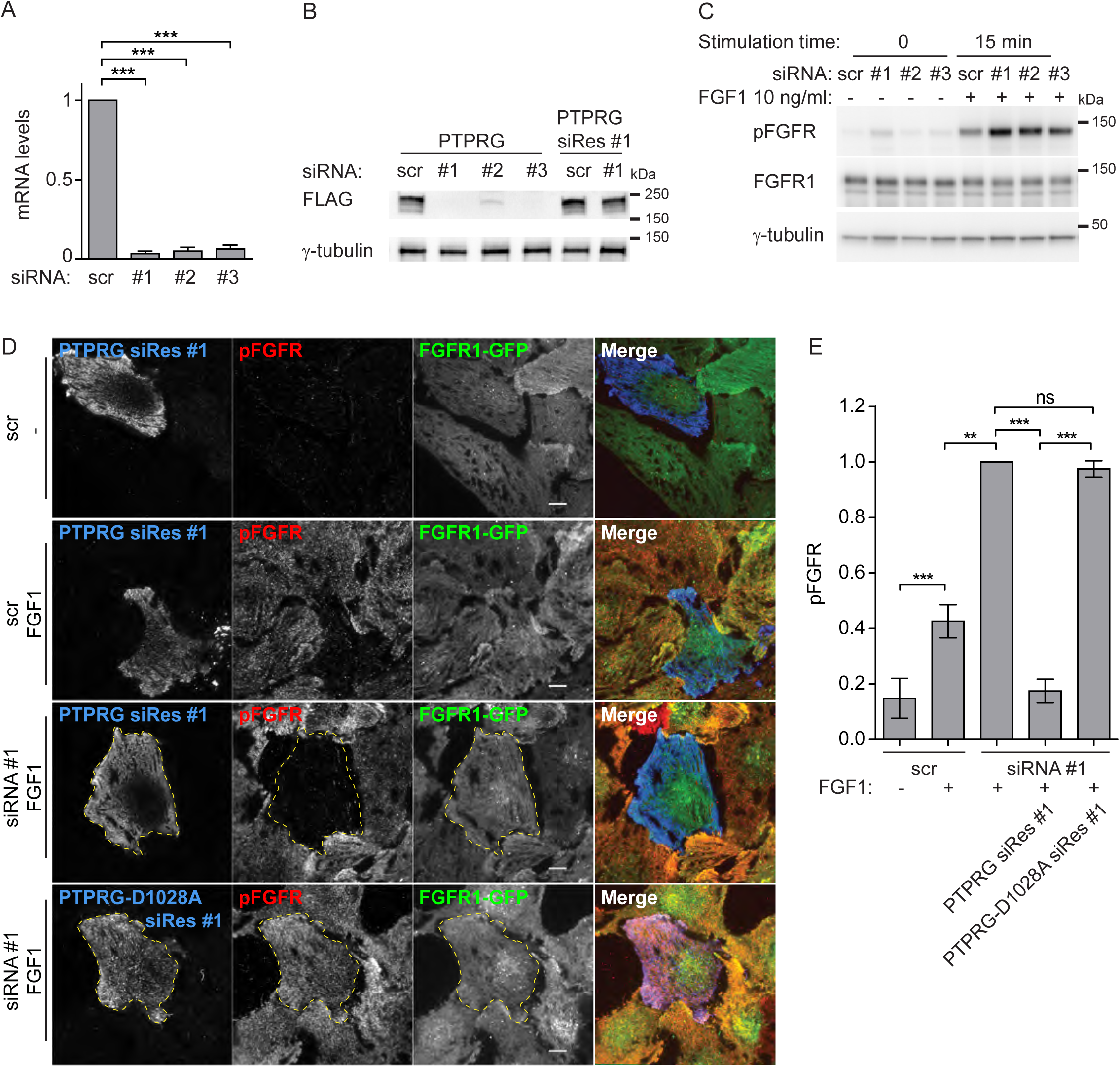
PTPRG knockdown increases FGFR1 autophosphorylation. **(A)** U2OS-R1 cells transfected with indicated siRNAs for 72 hours were lysed and RNA isolation, cDNA synthesis and qRT-PCR were performed as described in materials and methods. The amount of mRNA was calculated relative to the housekeeping gene SDHA and is expressed as fraction of scr. The histograms represent the mean + SD of three independent experiments. ***p ≤ 0.001. **(B)** Cells transfected with indicated siRNAs for 18 hours were transfected with MYC-FLAG-tagged PTPRG or siRNA-Resistant PTPRG (PTPRG siRes #1). 24 hours later cells were lysed and the lysates were subjected to SDS-PAGE followed by western blotting using denoted antibodies. A representative western blot is shown. **(C)** U2OS-R1 cells were treated with PTPRG siRNAs (#1-#3) or control siRNA (scr) for 72 hours. The cells were then serum-starved for 2 hours before stimulation with 10 ng/ml FGF1 in the presence of 10 U/ml heparin for 15 minutes. Next, the cells were lysed and the lysates were subjected to SDS-PAGE followed by western blotting using denoted antibodies. A representative western blot is shown. **(D)** U2OS-R1-GFP cells were transfected with control (scr) or PTPRG-specific siRNA (siRNA #1) for a total of 72 hours, transfected with MYC-FLAG-tagged siRNA resistant PTPRG (PTPRG siRes #1) or PTPRG-D1028A (PTPRG-D1028A siRes #1), for 20 hours. The cells were then starved for 2 hours, and stimulated (or not) with FGF1 in the presence of heparin for 10 minutes and then fixed and stained with anti-FLAG, and anti-phospho-FGFR (pFGFR) and fluorophore labeled secondary antibodies. The cells were imaged by TIRF and representative cells are shown. Merged images are overlays of PTPRG in blue, pFGFR in red, and FGFR1-GFP in green. Scale bars 8 μm. **(E)** The signal intensities for pFGFR in PTPRG-transfected or -untransfected cells were measured for 15-30 cells for each condition in four independent experiments and is presented as the mean values ± SD where values had been normalized to the signal intensity of cells that were transfected with PTPRG-specific siRNA, but not expressing tagged PTPRG, and stimulated with FGF1. The data were analysed using one-way RM ANOVA followed by Tukey *post hoc* test. ***p ≤ 0.001, **p ≤ 0.01, ns - not-significant.

### PTPRG downregulates FGFR activation in osteosarcoma cells

To further confirm the regulation of FGFR1 autophosphorylation by PTPRG and to analyse how this impinges on down-stream signalling pathways in osteosarcoma cells, we depleted cells for PTPRG using three different siRNAs and probed the activation of FGFR1 and its downstream signalling pathways by western blotting using phospho-specific antibodies (Fig. 5).

**Figure 5.**
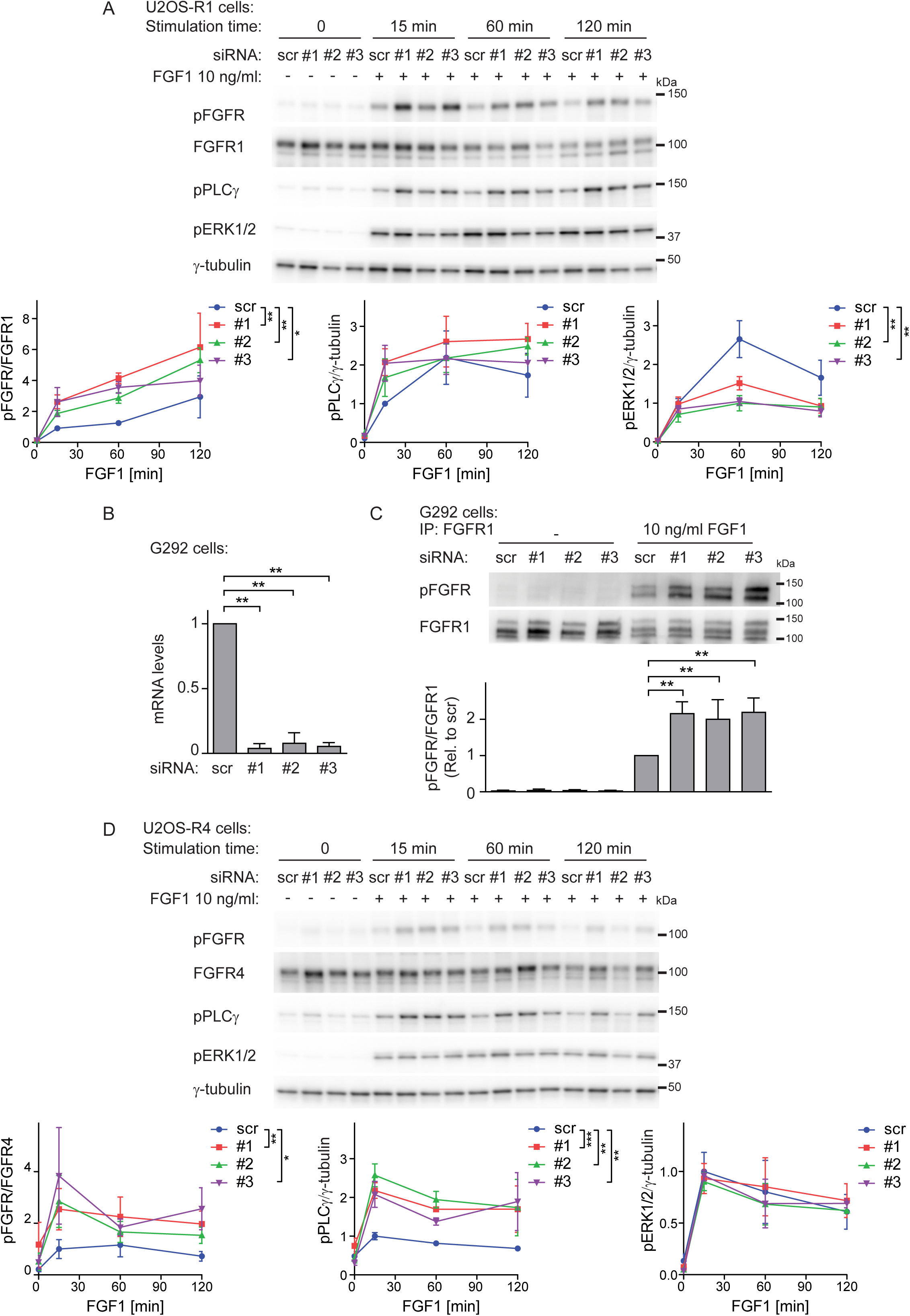
Increased FGFR1 signalling upon PTPRG knockdown. **(A)** U2OS-R1 cells were treated with PTPRG siRNAs (#1-#3) or control siRNA (scr) for 72 hours. Then the cells were serum-starved for 2 hours followed by stimulation with 10 ng/ml FGF1 in the presence of 10 U/ml heparin for various time points. The cells were then lysed and the lysates were subjected to SDS-PAGE followed by western blotting using denoted antibodies. The western blots were quantified and bands corresponding to phosphorylated proteins were normalized to total FGFR1 or loading control (γ-tubulin) (as indicated) and presented as fraction of scr, at 15 minutes stimulation time point. Means ± SEM of three independent experiments are presented on the graphs. The time-course series were analysed together using two-way ANOVA. ***p ≤ 0.001, **p ≤ 0.01, *p ≤ 0.05. **(B)** G292 cells transfected with indicated siRNAs for 72 hours were lysed and RNA isolation, cDNA synthesis and qRT-PCR were performed as described in materials and methods. The amount of mRNA was calculated relative to the housekeeping gene SDHA and is expressed as fraction of scr. The histograms represent the mean + SD of three independent experiments. **p ≤ 0.01. **(C)** G292 cells were left untreated or treated with 10 ng/ml FGF1 in the presence of 10 U/ml heparin for 20 minutes. The cells were then lysed and the cell lysates were subjected to immunoprecipitation using anti-FGFR1 antibody followed by SDS-PAGE and western blotting with indicated antibodies. Quantification of western blots are shown below. Bands corresponding to phosphorylated FGFR1 (pFGFR) were normalized to total FGFR1. The graph represents the mean + SD of three independent experiments. **p ≤ 0.01. **(D)** U2OS-R4 cells were treated as in **(A)**, lysed and the lysates were subjected to SDS-PAGE followed by western blotting using denoted antibodies. The western blots were quantified and bands corresponding to phosphorylated proteins were normalized to total FGFR1 or loading control (γ-tubulin) (as indicated) and presented as fraction of scr, at 15 minutes stimulation time point. Means ± SEM of three independent experiments are presented in the graphs. The time-course series were analysed together using two-way ANOVA. ***p ≤ 0.001, **p ≤ 0.01, *p ≤ 0.05.

U2OS-R1 cells depleted of PTPRG displayed significantly increased levels of phosphorylated FGFR1 upon FGF1 stimulation for 15-120 minutes compared to control cells (Fig. 5A). Interestingly, also the down-stream signalling molecule PLCγ displayed increased activity when PTPRG was depleted. However, in the case of ERK1/2 activation, we observed no increase after 15 minutes of FGF1 treatment and even a decrease during the later time points.

We also evaluated the effect of PTPRG depletion in the osteosarcoma cell line G292, expressing endogenous FGFR1. Efficient knockdown was confirmed by realtime PCR (Fig 5B). Since the expression level of FGFR1 is relatively low in this cell line, we immunoprecipitated the receptor using anti-FGFR1 antibodies and protein G coupled beads before analysis of phosphorylation levels by western blotting. Upon 15 minutes of stimulation with FGF1 we observed increased activation of FGFR1 in PTPRG depleted cells (Fig. 5C), which shows that PTPRG can downregulate endogenous FGFR1 and confirms the previous findings.

To test if PTPRG also regulates other FGFR family members, we performed experiments using cell lines stably expressing FGFR4 or FGFR2. We observed upregulated autophosphorylation of FGFR4 during 15-120 minutes stimulation by FGF1 in U2OS-R4 cells depleted of PTPRG, and a parallel increase of PLCγ phosphorylation (Fig. 5D), whereas no change was observed for ERK1/2 activation (Fig. 5D). PTPRG also regulated FGFR4 autophosphorylation in the rhabdomyosarcoma cell line RH30 expressing endogenous FGFR4 (Fig. 5-supplement 1A). Also in U2OS cells stably expressing FGFR2, we observed a similar increase in autophosphorylation of the receptor upon depletion of PTPRG (Fig. 5- supplement 1B). The results confirm that PTPRG down-regulates autophosphorylation of several FGFR family members and that an excessive activation of FGFRs, due to the loss of PTPRG, can lead to an elevation of downstream signalling pathways.

### PTPRG regulates the biological response to FGF1

Since our results demonstrate that PTPRG is responsible for dephosphorylation of activated FGFR, we hypothesized that the phosphatase could possibly alter the balance between phosphorylated and unphosphorylated forms of ligand-bound receptors. To confirm this hypothesis we evaluated the levels of phospho-FGFR1 in PTPRG-depleted osteosarcoma cells stimulated with various concentrations of FGF1.

First, we evaluated whether depletion of PTPRG alters the sensitivity of FGFR1 towards FGF1 stimulation. After 15 minutes of treatment with 0-20 ng/ml FGF1, the levels of FGFR1 phosphorylation and activation of downstream signalling pathways were analysed by western blotting. We observed increased activation of FGFR1 and PLCγ upon PTPRG knockdown under the applied range of FGF1 stimulation (Fig. 6A). No changes in ERK1/2 activation were observed. A detailed quantification of phospho-FGFR1 bands enabled us to detect a significant shift in the FGF1 dose-response curve towards higher pFGFR1 values when PTPRG was depleted (Fig. 6B). This demonstrates that PTPRG decreases the sensitivity of FGFR1 activation in response to FGF1. The effect of PTPRG depletion was more pronounced in parallel with increasing FGF1 concentrations. This agrees with the hypothesis that FGFR1 is a substrate for PTPRG.

**Figure 6.**
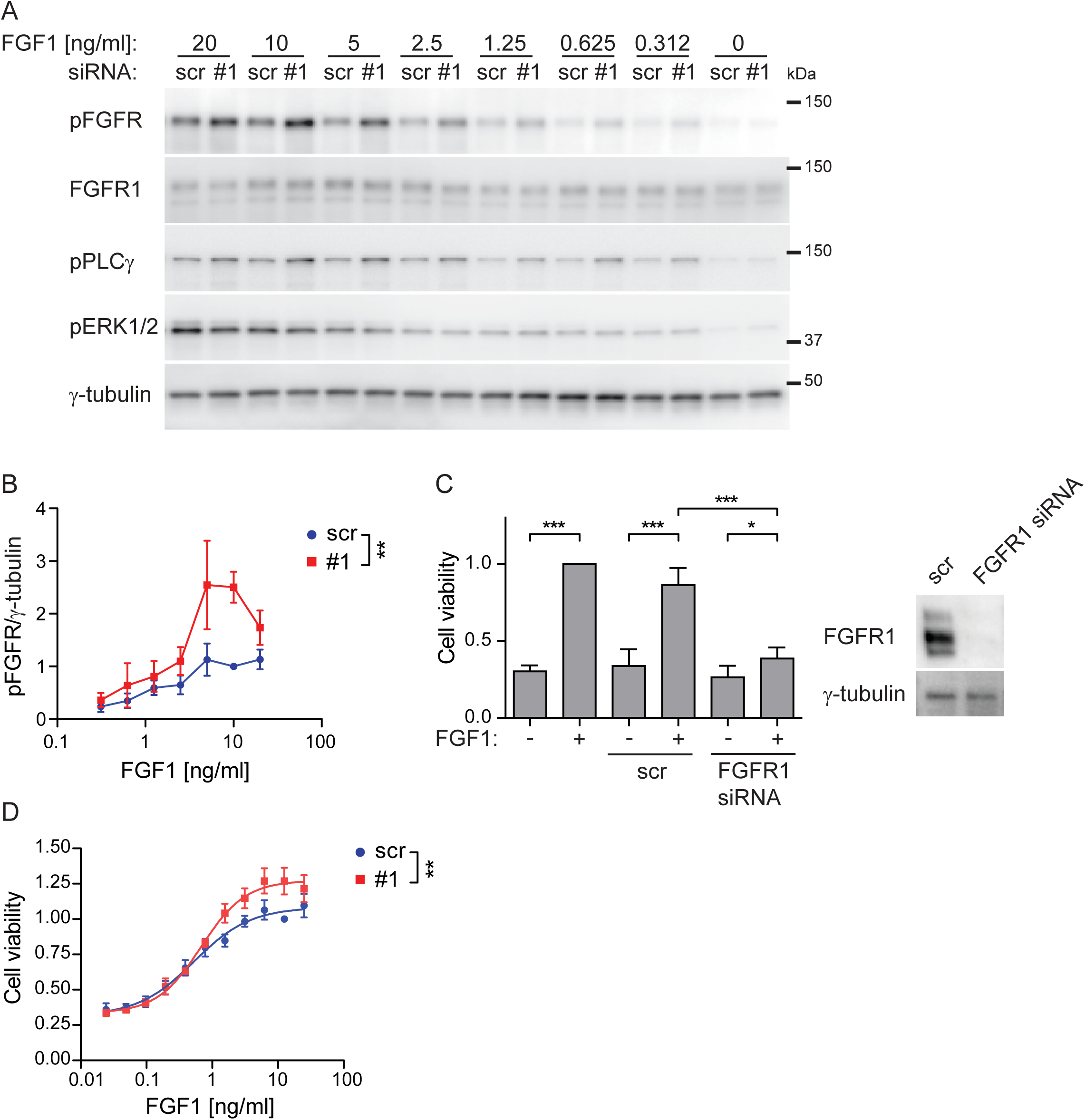
PTPRG regulates cellular sensitivity to FGF1. **(A)** U2OS-R1 cells were treated with siRNAs against PTPRG (#1) or control siRNA (scr) for 72 hours. Then the cells were serum-starved for 2 hours and stimulated with various concentrations of FGF1 in the presence of 10 U/ml heparin for 15 minutes, lysed and the lysates were subjected to SDS-PAGE followed by western blotting using denoted antibodies. A representative western blot is shown. **(B)** Western blots were quantified and bands corresponding to phosphorylated FGFR1 (pFGFR) were normalized to loading control (γ-tubulin) and presented as fraction of scr, 10 ng/ml stimulation. The graph represents the mean ± SEM of three independent experiments. The concentration series were analysed together using two-way ANOVA. **p ≤ 0.01. **(C)** G292 cells were left untreated or treated with FGFR1 siRNA or control siRNA (scr) for 72 h. During the last 48 hours the cells were treated with 100 ng/ml FGF1 in the presence of 10 U/ml heparin. Cell viability was then measured using CellTiter-Glo assay. The obtained data were normalized to non-transfected cells, stimulated with FGF1. Three technical replicates for each condition were included in each experiment. The graph represents the mean + SD of four independent experiments. ***p ≤ 0.001, *p ≤ 0.05. A representative western blot showing the knockdown efficiency of FGFR1 after 72 hours are presented to the right. **(D)** G292 cells were treated with PTPRG siRNA (#1) or control siRNA (scr) for 72 hours and then stimulated with different concentrations of FGF1 in the presence of 10 U/ml heparin for 48 hours. Cell viability was then measured using CellTiter-Glo assay. Four technical replicates for each condition were included in each experiment. The obtained data were normalized to scr, 12.5 ng/ml FGF1 and presented in the graph as means ± SEM of 4 independent experiments. The fitted curve represents non-linear regression analysis using Hill equation (dose-response with variable slope). The concentration series were analysed together using two-way ANOVA. **p ≤ 0.01.

We also tested whether the increased sensitivity of PTPRG-depleted cells towards FGF1 is biologically relevant. We chose the G292 cell line, expressing endogenous levels of FGFR1, and which growth in serum free media is dependent on FGF1 (Fig. 6C). We found that PTPRG-depleted cells displayed increased viability after treatment with various concentrations of FGF1 for 48 hours (Fig. 6D). We also found that the difference was more pronounced with increasing concentrations of FGF1, in correspondence with the results obtained for analysis of FGFR1 phosphorylation by western blotting (Fig. 6B). Our data indicate that PTPRG restricts the efficiency of the biological response of cells to FGF, and moreover, that down-regulation of PTPRG can serve as an advantage for cancer cells expressing FGFR1, allowing them to respond to lower FGF levels.

### Altered drug sensitivity in cells depleted for PTPRG

Given that PTPRG counter the activity of FGFR by dephosphorylation, we wanted to test if PTPRG could influence the action of a small molecule tyrosine kinase inhibitor on FGFR1. Since our data suggest that PTPRG is involved in shifting the balance of receptor autophosphorylation to the inactive, non-phosphorylated state, the phosphatase could also affect kinase inhibition.

We first tested increasing concentrations of the FGFR kinase inhibitor AZD4547 on U2OS-R1 cells stimulated with a constant amount of FGF1 (10 ng/ml) and investigated the levels of activated FGFR1 and its downstream signalling molecules by western blotting. We found that PTPRG-depleted cells displayed a higher level of phosphorylated FGFR1 and PLCγ in the presence of the FGFR kinase inhibitor (Fig. 7A). Little effect was observed on ERK activation. Quantification of phospho-FGFR1 bands visualized a shift in the dose-response curve when PTPRG was knocked down (Fig. 7B). These data indicate that higher concentrations of the inhibitor are needed to prevent FGFR1 kinase activity when PTPRG is downregulated.

**Figure 7.**
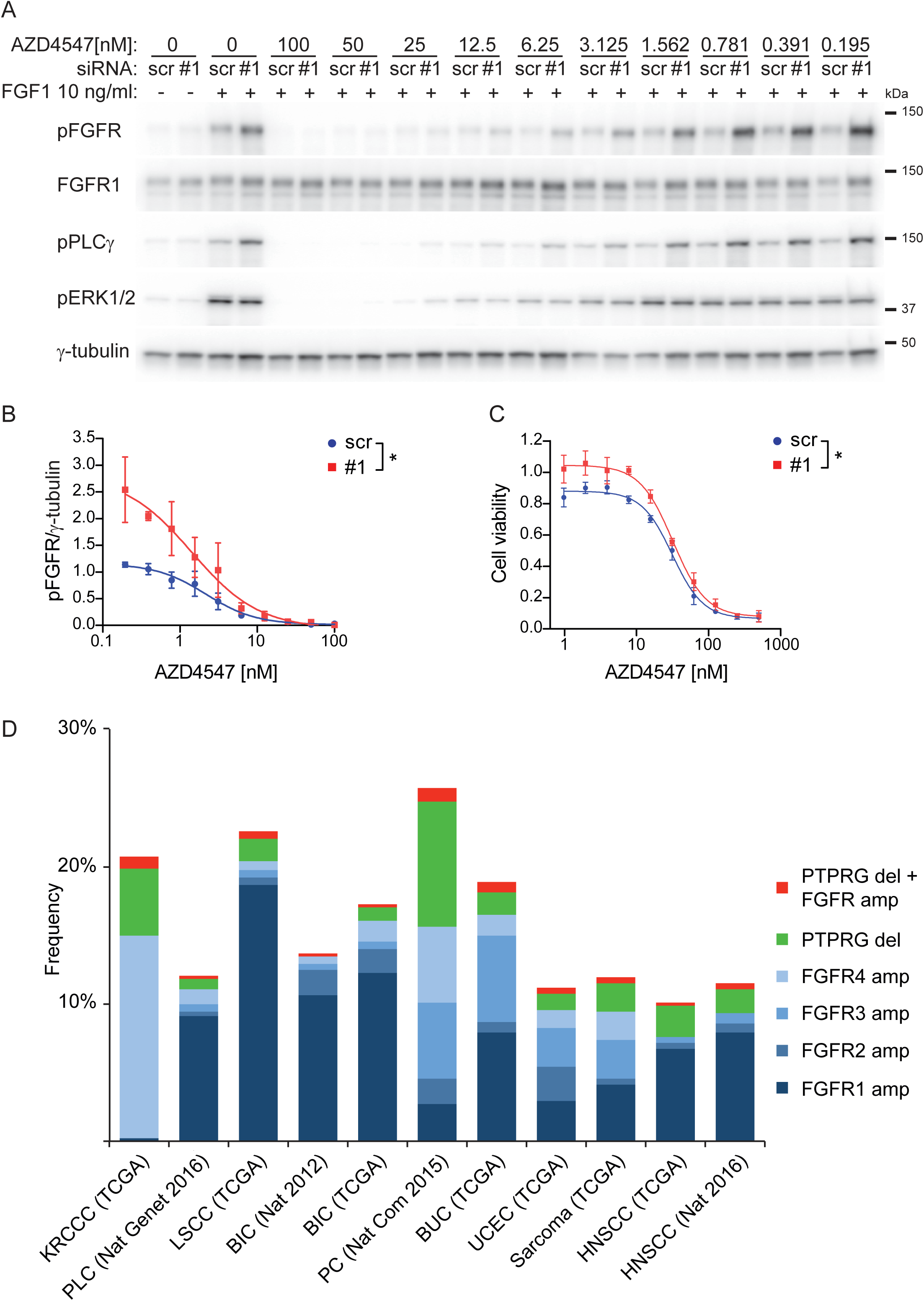
PTPRG influences the efficiency of FGFR inhibitors. **(A)** U2OS-R1 cells were treated with siRNAs against PTPRG (#1) or control siRNA (scr) for 72 hours, serum-starved for 2 hours and stimulated with 10 ng/ml FGF1 in the presence of 10 U/ml heparin and various concentrations of AZD4547 for 15 minutes. The cells were then lysed and the lysates were subjected to SDS-PAGE followed by western blotting using denoted antibodies. A representative western blot is shown. **(B)** Western blots were quantified and bands corresponding to phosphorylated FGFR1 (pFGFR) were normalized to loading control (γ-tubulin) and presented as fraction of scr, 10 ng/ml stimulation. The graph represents the mean ± SEM of three independent experiments. The fitted curve represents non-linear regression analysis using Hill equation (dose-response with variable slope). The inhibitor concentration series were analysed together using two-way ANOVA. *p ≤ 0.05. **(C)** G292 cells were treated with PTPRG siRNA (#1) or control siRNA (scr) for 72 hours and then stimulated with 10 ng/ml FGF1 in the presence of 10 U/ml heparin and various concentrations of AZD4547 for 48 hours. Cell viability was then measured using CellTiter-Glo assay. Four technical replicates for each condition were included in each experiment. The obtained data were normalized to scr without inhibitors and presented in the graph as means ± SEM of three independent experiments. The fitted curve represents non-linear regression analysis using Hill equation (dose-response with variable slope). The inhibitor concentration series were analysed together using two-way ANOVA. *p ≤ 0.05. **(D)** The graph shows the frequency of amplifications of the different FGFRs, deletions of PTPRG and the frequency of cases where at least one receptor is amplified and PTPRG is deleted. The figure is based on data generated by the TCGA Research Network (http://cancergenome.nih.gov/). The frequency is calculated according to the total number of cases for each study: KRCCC (Kidney Renal Clear Cell Carcinoma, 448 cases), PLC (Pan-Lung Cancer, 1144 cases), LSCC (Lung squamous cell carcinoma, 504 cases), BIC (Breast Invasive Carcinoma, 482 cases and 1105 cases), PC (Pancreatic Cancer, 109 cases), BUC (Bladder Urothelial Carcinoma, 127 cases), UCEC (Uterine Corpus Endometrial Carcinoma, 242 cases), Sarcoma (243 cases), HNSCC (Head and Neck Squamous Cell Carcinoma, 504 and 279 cases).

Next, we tested whether the disturbance in FGFR kinase inhibition, as a result of depleted PTPRG, translates into efficiency of the inhibitor to decrease cell growth. We knocked down PTPRG in G292 cells before stimulation with 10 ng/ml FGF1 in the presence of various concentrations of AZD4547. The experiment was performed using serum-free media to allow the cell growth to be dependent solely on FGF1. We found that PTPRG-depleted cells exhibited elevated viability after 48 hours of FGF1 treatment in the presence of AZD4547 (Fig. 7C). The difference was dependent on the concentration of the inhibitor, being more pronounced at lower concentrations. Our findings suggest that higher concentrations of FGFR inhibitors are necessary to control FGFR activity in cells with low levels of PTPRG. Importantly, this effect would imply an advantage for cancer cells lacking PTPRG and serve as a possible resistance-mechanism to FGFR inhibitors.

To explore among different cancer types the frequency of cases where at least one FGFR is amplified and PTPRG is deleted, TCGA data generated by the TCGA Research Network (http://cancergenome.nih.gov/) was investigated. Specific gene information was extracted from 11 studies showing frequent FGFR amplifications by using the cBioportal (http://www.cbioportal.org/) (Gao et al., 2013, Cerami et al., 2012). The results show that the frequency of cases with amplified receptor and deleted PTPRG is between 0.2-0.9% and is frequently found across many different cancer types (Fig. 7D). In total, 18 cases of the combination FGFR amplification/PTPRG deletion was identified in the 11 studies investigated, clearly demonstrating the relevance of our findings in human cancer.

## DISCUSSION

FGFR inhibitors are now entering the clinic and it is crucial to understand how tumour cells respond to this treatment (Dieci et al., 2013). We show here that PTPRG, a membrane bound tyrosine phosphatase, is an important modulator of FGFR tyrosine kinase activity. We demonstrate that PTPRG counter the activity of FGFR1 by direct dephosphorylation of the autoactivated, tyrosine phosphorylated FGFR1. The activity of PTPRG is also a determinant of the efficacy of FGFR inhibitors. We found that lowering the levels of PTPRG by specific siRNAs protected tumour cells against the clinically relevant FGFR kinase inhibitor AZD4547. It is therefore possible that PTPRG levels in cancer cells could be a predictor of outcome of FGFR kinase inhibition. Our data suggest that in clinical trials using FGFR inhibitors the level of PTPRG should be determined, in order to test the possibility that in tumours with low levels of PTPRG, kinase inhibitors may not be as efficient as in cells with normal levels of PTPRG. This may be particularly important when treating tumours with low doses of kinase inhibitors, which is normally the case since these inhibitors are associated with severe toxicity (e.g. hyperphosphatemia and tissue calcification) (Dieci et al., 2013).

Interestingly, we observed a difference in the response of two downstream signalling pathways to PTPRG depletion. While the activity of PLCγ, similarly to that of FGFR, was upregulated, ERK phosphorylation was mainly unchanged, and even reduced at later time points. The reason for this phenomenon could be that the MAPK pathway is subjected to several layers of both positive and negative regulation that may buffer for increased activity of the receptor. This may also imply that viability in osteosarcoma cells is modulated by other signalling pathways than the MAPK pathway.

Our studies suggest that PTPRG’s main cellular localization is at the cell surface and that it is inefficiently endocytosed compared to the FGF1/FGFR1 activated complex. Concurrently, we find that PTPRG levels profoundly affect FGFR1 activity at the early timepoints (minutes) after FGF1 stimulation, which initiates at the plasma membrane. We also find that PTPRG levels affect FGFR1 and downstream signalling events even 2 hours after the initial stimulation, and that this translates into biological effects such as cell viability several days after onset of FGF1 stimulation. It is not known in detail how the FGFR1 activity is affected by its subcellular localization, i.e. whether the rate of FGFR1 autophosphorylation is maintained or reduced after transfer from the plasma membrane to the endosomal membrane. Possibly, PTPRG levels exert a long term effect on FGFR1 activity mainly by regulating its initial activation rate.

Sarcoma cells were here used to study the regulation of FGFR1, but it is likely that PTPRG also dephosphorylates FGFR1 in other cell types. For example, FGFR1 is overexpressed in breast cancers and is an attractive target with several clinical trials under way. Interestingly, TCGA data show that PTPRG is deleted and mutated in a subset of breast cancer patients (Fig 7D). Intriguingly, there are also reported cases where FGFR1 is overexpressed and PTPRG deleted, which could possibly be a particular bad combination for the patient. We also show that FGFR1 becomes hypersensitive to its ligand when PTPRG is down-regulated. It is therefore possible that FGFR1 can be aberrantly activated by low levels of ligand in the tumour microenvironment causing tumorigenic growth without overexpression or mutation of the receptor itself.

We show in this study that also other FGFR members (FGFR2 and FGFR4) are regulated by PTPRG. FGFR2 is activated by mutations and is an attractive target in endometrial cancer, and FGFR4 is a potential drug target in rhabdomyosarcoma. The identification of PTPRG as a potent regulator of FGFR activity may therefore have broad consequences in cancer therapy.

We used BioID to investigate proximal proteins to FGFR1 that could potentially regulate FGFR signalling. The advantage of this method is that the biotinylation occurs in living cells and that the biotin tag makes it easier to pull down transient interactors and transmembrane proteins that may be difficult to detect in classic pull-down assays (Roux et al., 2013). Indeed, PTPRG has not been found in any previous studies where FGFRs have been co-precipitated. Thus, as shown here, BioID may be used to find important interactors that have not been found with the standard methods.

We have here concentrated our efforts on PTPRG, but we believe our proteomic data may be a resource for further studies of the regulation of FGFR signalling. For instance, in FGF1-stimulated cells, we identified known downstream effectors of activated FGFR (e.g. PLCγ, RSK2 and SHC4), but we also uncovered members of other signalling pathways (Fig. 1 supplement 4 and Table S1). For instance, we found several members of the interferon-stimulated gene family, which may play a role in immunity. We also identified two cyclins (CCNE1, CCNB2) suggesting that FGFR1 may interact directly with these cell cycle regulators to stimulate proliferation. As we also have shown recently for FGFR4(Haugsten et al., 2016), BioID revealed association with the FGFRs and a number of proteins involved in vesicular trafficking, reflecting the importance of intracellular transport for these receptors. Analysing proteins whose expression was induced by FGF1 signalling, we found several proteins that may confer negative feedback (Table S1). Examples include A2M, which has previously been shown to bind the ligand FGF2 and thereby blocking its interaction with the receptors and the heparan sulfate proteoglycan CD44 that has been shown to regulate FGFR action. We also noticed a protein that has been shown to be a feedback inhibitor for EGFR family members (ERRFI1), which may possibly play a similar role for FGFRs. Finally, we also identified a phosphate transporter (SLC20A1) that may be involved in the reabsorption of phosphate mediated by FGFR signalling (Prie and Friedlander, 2010). This may indicate a more direct activation of phosphate transporters than previously anticipated.

It is known that PTPRG has other targets than FGFR1 (Cheung et al., 2015), but it remains an interesting question if additional tyrosine phosphatases are involved in directly regulating the activity of FGFR1. Indeed, two additional tyrosine phosphatases were discovered through our screen, while only PTPRG was among the top hits. However, the very strong effect of PTPRG knockdown on FGFR activity observed in our studies, indicates that PTPRG is a major regulator of FGFR, and also indicates that there may be less redundancy among phosphatases than anticipated. This also implies that cells with low expression of PTPRG may be particularly vulnerable to excessive FGFR activity, which could lead to more aggressive cancer. We therefore believe that it will be important to study PTPRG as a predictor of outcome for disease caused by FGFRs.

## Methods

### Antibodies and compounds

The following antibodies were used: rabbit anti-FGFR1 (ab76464), and rabbit anti-Clathrin heavy chain (ab21679) from Abcam; rabbit anti-FGFR1 (2144-1) from Epitomics; mouse anti-phospho-FGFR (Tyr653/654) (#3476), rabbit anti-DYKDDDDK (FLAG) tag (#2368), rabbit anti-phospho-PLCγ (Tyr783) (#14008), mouse anti-phospho-ERK1/2 (Thr202/Tyr204) (#9106) from Cell Signaling Technology; mouse anti-γ-tubulin (T6557), and mouse anti-FLAG M2 antibody (F-1804) from Sigma-Aldrich; mouse anti-EEA1 (610456) from BD transduction laboratories; rabbit anti-phospho-PLCγ (Tyr783) (sc-12943-R) from Santa Cruz Biotechnology; rabbit anti-HA epitope tag (600-401-384) from Rockland; mouse anti-MYC Tag (05-724) from Merck Millipore, human anti-EEA1 antiserum was a gift from B. H. Toh (Monash University), HRP-Streptavidin (016-030-084), Alexa488-Streptavidin (016-540-084) and all secondary antibodies from Jackson Immuno-Research Laboratories.

Protease inhibitor cocktail tablets (ethylenediaminetetraacetic acid (EDTA)-free, complete) were from Roche Diagnostics. DyLight 550 NHS Ester, Ez-link Sulfo-NHS-SS-biotin, Pierce^™^ anti-HA magnetic beads and Dynabeads G protein were from Thermo Scientific. Hoechst 33342 was purchased from Life Technologies. Streptavidin Sepharose High Performance was from GE Healthcare Life Sciences. Mowiol, biotin, heparin, PD173074, active human PTPRG catalytic domain (SRP0223), active human PTPN12 catalytic domain (SRP5073), p-nitrophenyl phosphate (pNPP) and phosphatase inhibitors were from Sigma-Aldrich. AZD4547 was purchased from SelleckChem. FGF1 was prepared as previously described (Wesche et al., 2005). FGF1 was labeled with DyLight 550 following the manufacturer's procedures. Recombinant GST, expressed in *E. coli* and purified using GSH Sepharose Fast Flow (GE Healthcare), was kindly provided by Dr. Coen Campsteijn from the Department of Molecular Cell Biology, Institute for Cancer Research, Oslo University Hospital.

### Plasmids and siRNAs

pcDNA3.1-FGFR1-BirA* was made by cloning a PCR fragment containing the FGFR1 open reading frame and AgeI-HF and BamHI-HF flanking sites into pcDNA3.1 MCS-BirA*(R118G)-HA cut with AgeI-HF and BamHI-HF using pcDNA3-hFGFR1 as a template (Haugsten et al., 2005). pcDNA3.1 MCS-BirA(R118G)-HA was a gift from Kyle Roux (Addgene plasmid # 36047) (Roux et al., 2012). Construction of the pcDNA3.1/Zeo-BirA* was described previously (Haugsten et al., 2016). pEGFP-FGFR1 was made by cloning a PCR fragment containing the FGFR1 open reading frame and XhoI and ApaI flanking sites into pEGFP-N1 cut with XhoI and ApaI using pcDNA3-hFGFR1 as template. pCMV6-Entry vector containing PTPRG-MYC-FLAG was purchased from Origene (RC_218964). PTPRG mutants were produced by site-directed mutagenesis using Pfu I HF (Agilent) with specific primers, followed by DpnI (New England Biolabs) treatment. PTPRG inactivating mutation (D1028A) was introduced with the primer 5’-TACACAGTGGCCTGCCATGGGAGTTCCCG-3’, while the primer 5’-CATTAGCCATGTCTCACCCGATAGTCTATATTTATTTCGGGTCCAGGCCGTGTGTCGGAACGAC-3’ was used to mutate 7 nucleotides and obtain siRNA-Resistant PTPRG (siRes#1 PTPRG) in both wild-type and D1028A mutant PTPRG. D1028A and siRes mutants were verified by sequencing. These plasmids are resistant to siRNA oligo s11549 (#1) PTPRG Silencer^®^ Select. Silencer^®^ Select siRNA oligos targeting PTPRG; s11549 (#1), s11550 (#2) and s11551 (#3), siRNA oligos targeting FGFR1 (s5177) and *Silencer^®^* select Negative Control No. 2 siRNA (scr) (4390846) were purchased from Life Technologies.

### Cells and transfection

To obtain U2OS cells stably expressing FGFR1-BirA* (U2OS-R1-BirA*), FGFR1-GFP (U2OS-R1-GFP) and FGFR2 (U2OS-R2) and U2OS-R1 stably expressing BirA* (U2OS-R1 + BirA*), Fugene liposomal transfection reagent was used according to the manufacturer’s protocol. Clones were selected with 1 mg/ml geneticin (U2OS-R1-BirA*, U2OS-R1-GFP and U2OS-R2) or 0.2 mg/ml Zeocin (U2OS-R1 + BirA*). Clones were chosen based on their receptor/BirA* expression levels analysed by immunofluorescence and western blotting. U2OS cells stably expressing FGFR1 has been described previously (Haugsten et al., 2008). The G292 and RH30 cell lines were generous gifts from Prof. Ola Myklebost (Department of Tumor Biology, The Norwegian Radium Hospital). U2OS and G292 cells were propagated in DMEM or RPMI (respectively) supplemented with 10% fetal bovine serum, 100 U/ml penicillin, and 100 μg/ml streptomycin in a 5% CO_2_ atmosphere at 37°C.

siRNA transfection was performed using Lipofectamine RNAiMAX Transfection Reagent (Invitrogen, Life Technologies) according to the manufacturer’s protocol. 10 nM of siRNA was used and the experiments were performed 72 hours after transfection. Transient expression of different plasmids was performed by transfecting cells with plasmid DNA using Fugene 6 Transfection Reagent (Roche Diagnostics) according to the manufacturer’s protocol.

### Mass Spectrometry

Affinity capture of biotinylated proteins, sample preparation and mass spectrometry was performed as previously described (Haugsten et al., 2016).

Experimental design and statistical rationale of the MS analysis: Six individual experiments were performed; three experiments consisting of samples C1 (U2OS-R1 cells), C2 (U2OS-R1 stably transfected with BirA*) and C3 (U2OS-R1 cells stably transfected with BirA* and stimulated with FGF1) and three experiments consisting of samples C1 (U2OS-R1 cells), S1 (U2OS-R1-BirA*) and S2 (U2OS-R1-BirA* stimulated with FGF1). All three samples in each of the six individual experiments were run three times (*n*=3 for LC variability, *n*=9 total number of replicates combined, in the case of C1: *n*=6 for LC variability, *n*=18 total number of replicates combined). In the case of one of the three experiments for C3 (U2OS-R1 cells stably transfected with BirA* and stimulated with FGF1) only one replicate was run (*n*=3 for LC viability, *n*=7 total number of replicates combined). The mean IBAQ values were calculated for each protein in each sample (C1, C2, C3, S1 and S2). Proteins identified in C1 were considered as background and the means of C3, S1 and S2 were compared to that of C1. Proteins were removed from the list if they were not significantly enriched at least ten times compared to C1 (p<0.05, two-tailed t test). Proteins identified in C2 were considered as BirA* background and the means of C3, S1 and S2 were next compared to that of C2. Proteins were removed from the list if they were not significantly enriched at least ten times compared to C2 (p<0.05, two-tailed t test). Proteins significantly enriched ten times or more in C3 compared to C1 and C2 were considered as proteins with potentially induced expression by FGF1 stimulation (p<0.05, two-tailed t test). Proteins significantly enriched ten times or more in S1 compared to C1 and C2 were considered as proteins in proximity to FGFR1. S2 was in addition to being compared to C1 and C2 also compared to C3 and proteins were removed from the list if they were not significantly enriched at least ten times compared to C3 (p<0.05, two-tailed t test). Proteins significantly enriched ten times or more in S2 compared to C1, C2, and C3 were considered as proteins in proximity to active FGFR1. The mass spectrometry proteomics data have been deposited to the ProteomeXchange Consortium via the PRIDE (Vizcaino et al., 2016) partner repository with the dataset identifier PXD006157. (Username, Password: 76TXQbfY)

### Western blotting

After indicated treatment, cells were lysed in lysis buffer supplemented with protease and phosphatase inhibitors or directly in sample buffer and the lysates were then loaded for SDS-PAGE (4-20% gradient) and afterwards transferred to a PVDF membrane (Bio-Rad) for western blotting. Blots were developed with SuperSignal West Dura Extended Chemiluminescent Substrate (Thermo Scientific) and detected using Gel Doc (Bio-Rad). Western blots were quantified using the Gel analysis function in Image J (Schneider et al., 2012).

### RNA isolation, cDNA synthesis and quantitative real-time polymerase reaction (qRT-PCR)

Total RNA was isolated from cell lysate using RNeasy plus minikit and the QIAcube robot (Qiagen) according to the manufacturer’s protocol. Then 0.5-1 mg of RNA was used for cDNA synthesis using iScript cDNA synthesis kit. Quantitative real-time PCR was performed using QuantiTect SYBR Green PCR kit, cDNA template and the following QuantiTect primers: PTPRG (QT00060116) and Succinate dehydrogenase (SDHA) (QT00059486). The qRT-PCR was run and analysed using the Lightcycler 480 (Roche). Cycling conditions were 5 minutes at 95°C followed by 45 cycles 10 seconds at 95°C, 20 seconds at 60°C and 10 seconds at 72°C. Gene amplification was normalized to the expression of SDHA.

### Co-immunoprecipitation assays

After indicated treatment, the cells were lysed in lysis buffer supplemented with protease and phosphatase inhibitors. Cell lysates were then subjected to immunprecipitation reactions with indicated antibody immobilized to Dyneabeads Protein G or with Pierce^™^ anti-HA magnetic beads. After washing, protein complexes were eluted in sample buffer, separated by SDS-PAGE and analysed by western blotting.

### *In vitro* phosphatase assay

The enzymatic activity of recombinant PTPRG (catalytic domain, residues 801-1147) and PTPN12 (catalytic domain, residues 1-355) was probed by a standard colorimetric assay using p-nitrophenyl phosphate (pNPP) as substrate (Lorenz, 2011). The initial reaction rate was monitored colorimetrically (Abs. at 405 nm) within the first 10 min of reaction, where the data fell in the linear range. The reaction buffer and 300 nM GST in reaction buffer served as control to exclude substrate self-degradation and the effect of potential impurities related to the GST fusion protein purification system. One unit of phosphatase activity (1 U) was defined as the amount of enzyme that hydrolyses 1 nmol of pNPP in 1 min at 30°C in 50 μl reaction volume. Molar extinction coefficient of the reaction product (pNP) was assumed as 18000 M^−1^cm^−1^.

After indicated treatment, U2OS-R1-BirA* cells were lysed in lysis buffer supplemented with protease and phosphatase inhibitors. Cell lysates were then subjected to immunprecipitation with Pierce^™^ anti-HA magnetic beads (Thermo Scientific), which were subsequently washed with lysis buffer without phosphatase inhibitors and incubated at 37°C with indicated recombinant phosphatases with addition of 2 mM DTT. The control samples were incubated with recombinant GST or in the presence of phosphatase inhibitor cocktail, as indicated in the figure legend. The immunoprecipitates were then eluted in sample buffer, separated by SDS-PAGE and analysed by western blotting.

### Light microscopy

For confocal microscopy, cells grown on coverslips were treated as indicated and fixed in 4% formaldehyde. The cells were then permeabilized with 0.1% triton X-100, stained with indicated antibodies and mounted in mowiol. Confocal images were acquired with a 63×objective on Zeiss LSM 780 and Zeiss LSM 710 confocal microscopes (Jena, Germany). Images were prepared with Zeiss LSM Image Browser and CorelDRAW11 (Ottawa, Canada).

For wide-field (WF) microscopy and structured illumination microscopy (SIM), U2OS-R1 cells were grown on 1.5H glass coverslips and transiently transfected with plasmid encoding MYC/FLAG-tagged PTPRG or PTPRG-D1028A using Fugene 6 (according to the producers procedures), for approx. 20 hours. The cells were serum starved for two hours (DMEM with penicillin and streptomycin but without serum), and then either fixed immediately or incubated with FGF1 (200 ng/ml) and heparin (10 U/ml) for 1 hour and then fixed.

For total internal reflection fluorescence (TIRF) microscopy, U2OS-R1-GFP cells were grown in glass bottom culture dishes (MatTek). The cells were transfected with plasmid encoding MYC/FLAG-tagged PTPRG or PTPRG-D1028A or siRNA resistant versions of these (using Fugene 6) for 20 hours. Next, the cells were serum-starved for 2 hours and stimulated for 10 min with FGF1 and heparin, FGF1 and heparin in the presence of PD173074 (including 30 min pretreatment with PD173074), or no FGF1, and then the cells were fixed. In some cases, the cells were also transfected with scrambled siRNA or siRNA against PTPRG (siRNA #1) two days prior to plasmid transfection.

Next, cells were fixed in 4 % formaldehyde (Sigma-Aldrich) in PBS (10 minutes at room temperature). The fixed cells were permeabilized with 0.05% saponin in PBS and stained with indicated combinations of primary antibodies diluted in PBS with 0.05% saponin, and anti-mouse/rabbit/human secondary antibodies labelled with Alexa Fluor 488, Alexa Fluor 568, or Alexa Fluor 647. Cells/coverslips for WF/SIM were also stained with Hoechst33342 and mounted on object slides with SlowFade Diamond Antifade Mountant (ThermoFisher). Stained cells for TIRF microscopy were maintained and imaged in PBS.

Wide-field, SIM, and TIRF imaging was performed on a Deltavision OMX V4 microscope (Applied Precision, Inc., Issaquah, WA) using an Olympus ×60 NA 1.42 Plan Apochromat objective for WF imaging and SIM, and an Olympus x60 NA 1.49 Plan Apo TIRF objective for TIRF imaging. The OMX is further equipped with an InSightSSI^™^ illumination module used for WF imaging, 405 nm, 488 nm, 568 nm, and 642 nm laserlines that were used for SIM and TIRF imaging, a Ring-TIRF module, and three cooled sCMOS cameras.

For WF imaging, z-stacks covering the whole cell were recorded with a z-spacing of 250 nm. For SIM, z-stacks were recorded with a z-spacing of 125 nm and for each focal plane, 15 raw images (five phases for three different angular orientations of the illumination pattern) were captured. WF images were deconvolved, SIM images were reconstructed, and all images were aligned using Softworx software (Applied Precision, Inc., Issaquah, WA).

All TIRF images were captured using the same channel specific settings for Ring-TIRF diameter, laser intensity and exposure. The phospho-FGFR1 signal was quantified using Fiji/ImageJ software as follows; Cells were selected for quantification based on GFP intensity (indicating average/normal FGFR1 levels), and identified as untransfected or transfected with PTPRG/PTPRG-D1028A based on FLAG-staining. ROI’s were defined by drawing the outline of selected cells, and the mean pixel value over an ROI in the phospho-FGFR1 specific channel was taken as the measure of the phospho-FGFR1 signal intensity of a cell. Images were subjected to background subtraction by a value set for each experiment. Data presented are the mean values of three or four independent experiments where 15-30 cells were measured for each condition in each experiment.

Further processing of images for presentations (projections, volume views, contrast adjustments, montages) were done using Fiji/ImageJ software.

### Cell viability assay

The cells were treated with indicated siRNAs and reseeded into 96-well plates the day before stimulation with FGF1 in serum-free medium, supplemented with 20 U/ml heparin,. The cells were treated with FGF1 72 hours after siRNA treatment. Cell viability was measured 48 h after stimulation using CellTiter-Glo assay (Promega). In the case of FGFR1 knockdown, cells were treated with FGFR1 siRNA or control siRNA (scr) for 72 hours. During the last 48 hours the cells were treated with 100 ng/ml FGF1 in the presence of 10 U/ml heparin. Cell viability was then measured using CellTiter-Glo assay.

### Statistical rationale

Data arised from series of three or more independent experiments as stated in figure legends. Results with *p*<*0.05* were considered statistically significant. Time-course and dose-response data series were analysed using two-way ANOVA. Single end-point assay data were analysed using one-way repeated measures (RM) ANOVA followed by Tukey *post hoc* test. For all experiments, the tests were performed on log transformed raw data. The tests were performed using GraphPad Prism 5 (GraphPad Software) or Sigma plot (Systat Software).

## Acknowledgments

We thank Dr. Coen Campsteijn, Prof. Ola Myklebost and Prof. Claudio Sorio for sharing reagents and Prof. Knut Liestøl for guidance on the statistics. This work was supported by the the Research Council of Norway through its Centers of Excellence funding scheme (project number 179571) and the Norwegian Cancer Society. J.W. holds a Researcher fellowship (project *5756681*), E.M.H a Career fellowship (project *6842225*), M.K. a Postdoctoral fellowship (project *734183*) and E.F. a Postdoctoral fellowship (project *5756681*) from the Norwegian Cancer Society. All MS data was collected at the Proteomics Core Facility, Rikshospitalet, which is funded by the Norwegian South-East Health Authority.

---

**Figure 1 – figure supplement 1.**
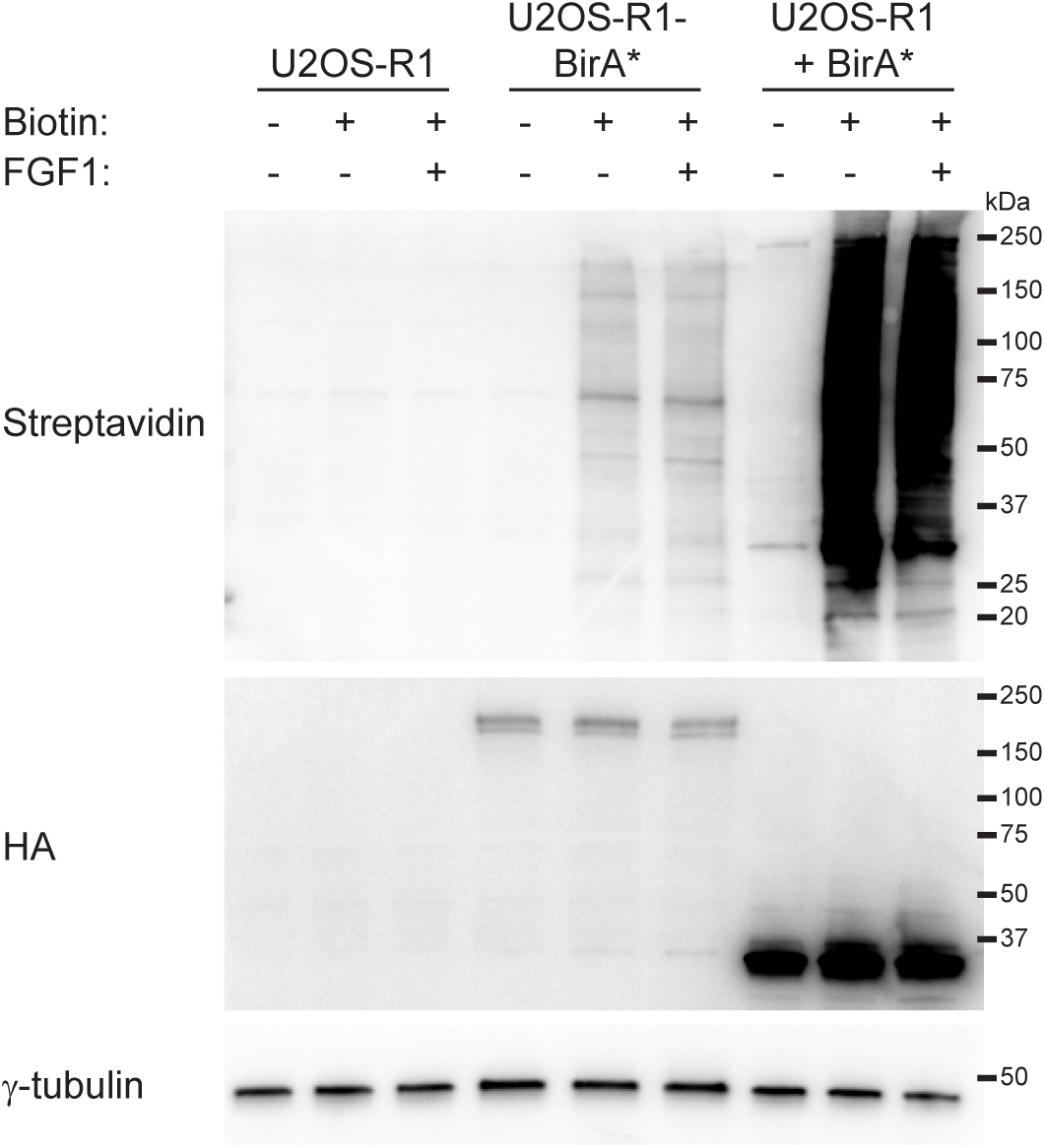
BioID of FGFR1 in osteosarcoma cells. U2OS-R1 cells, U2OS-R1-BirA* cells or U2OS-R1 cells coexpressing BirA* (U2OS-R1 + BirA*) were left untreated or treated with 50 mM biotin and/or 100 ng/ml FGF1 in the presence of 10 U/ml heparin as indicated for 24 hours. The cells were then lysed and the cellular material was analysed by SDS-PAGE and western blotting using the indicated antibodies. A representative western blot is shown.

**Figure 1 – figure supplement Table 1. Proteins identified by quantitative LC MS/MS.**

**Figure 1 – figure supplement 2.**
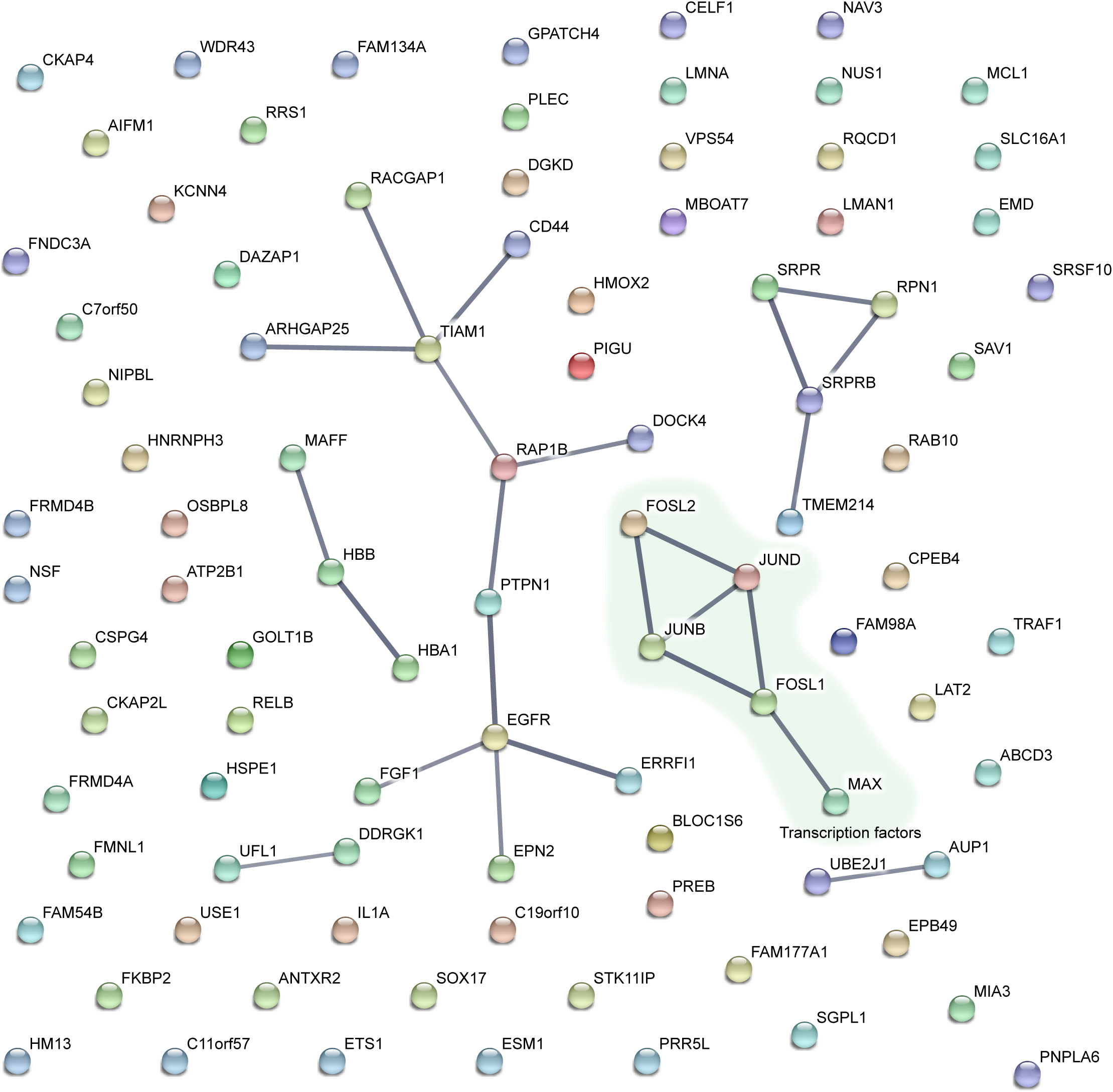
STRING interaction network. Version 10.0 of the STRING database was used (http://string-db.org/)(Szklarczyk et al., 2015) to investigate protein-protein interactions and construct an interaction network map for hits from the dataset of FGF1-induced expression (C3). Only known interactions from experiments and databases were included and a high confidence interaction score (> 0.7) was applied.

**Figure 1 – figure supplement 3.**
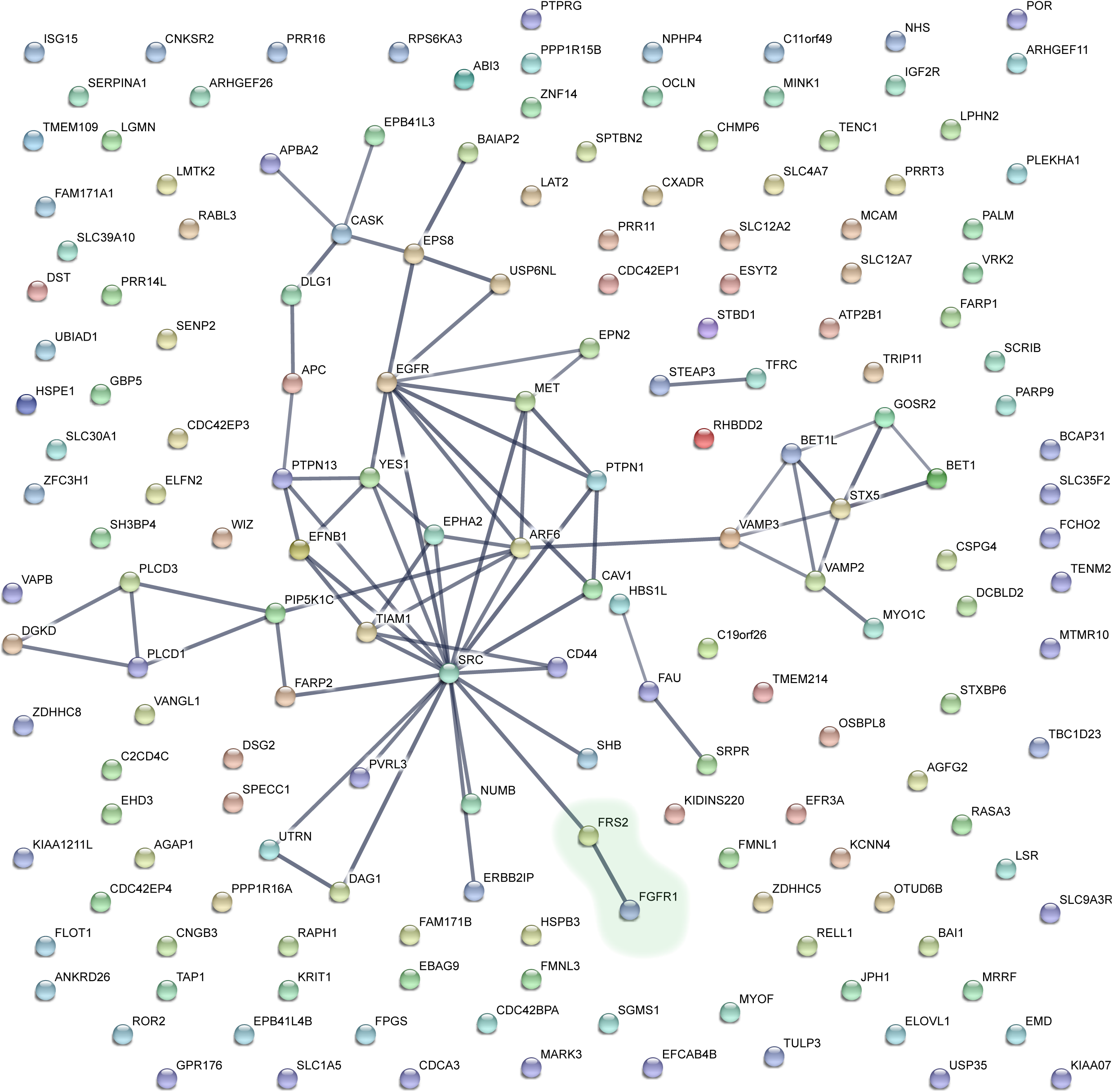
STRING interaction network. The dataset for proteins associated with unstimulated FGFR1 (S1) was subjected to the same analysis as in Fig. 1-figure supplement 2.

**Figure 1 – figure supplement 4.**
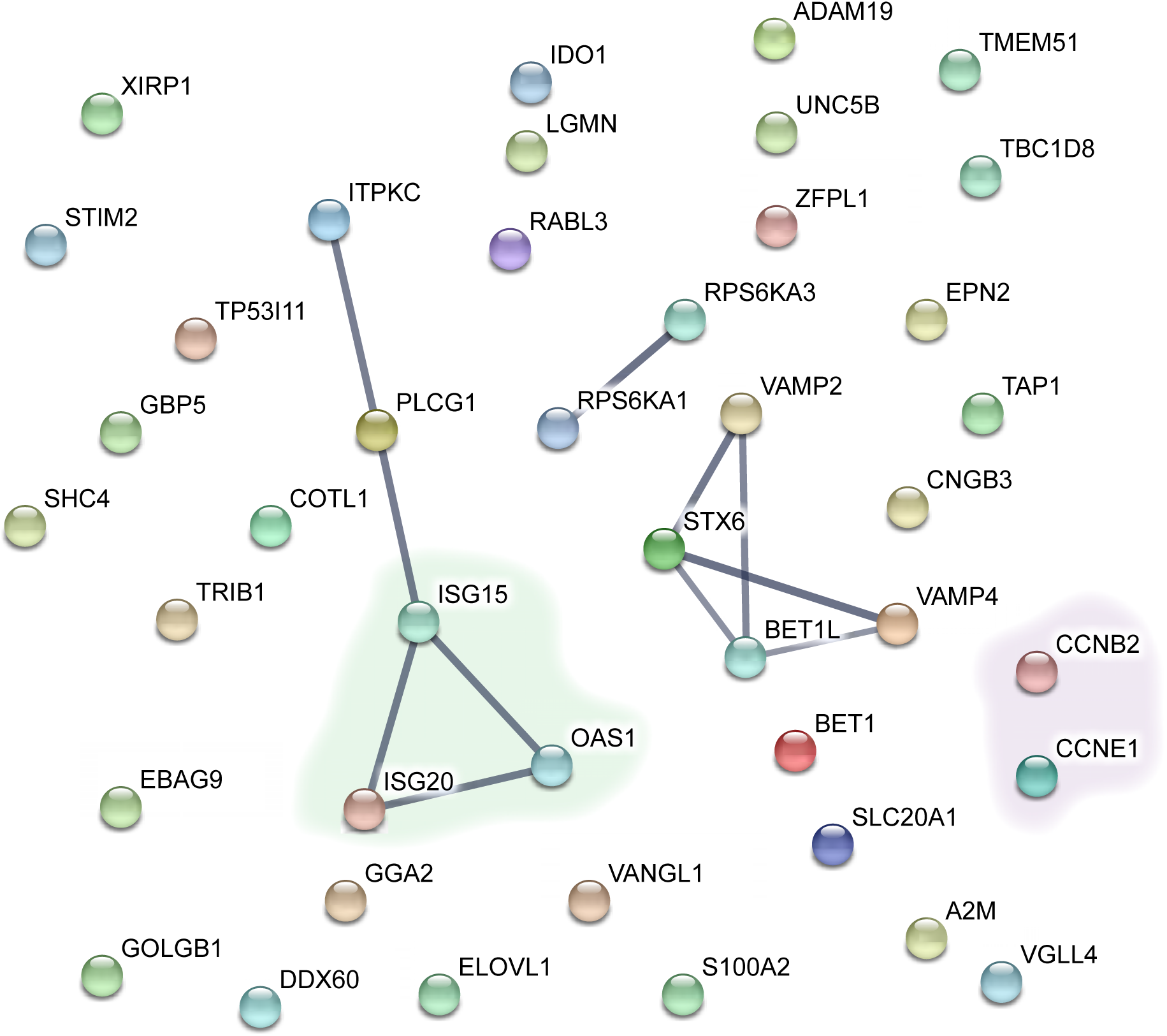
STRING interaction network. The dataset for proteins associated with FGF1-stimulated FGFR1 (S2) was subjected to the same analysis as in Fig. 1-figure supplement 2.

**Figure 5 – figure supplement 1.**
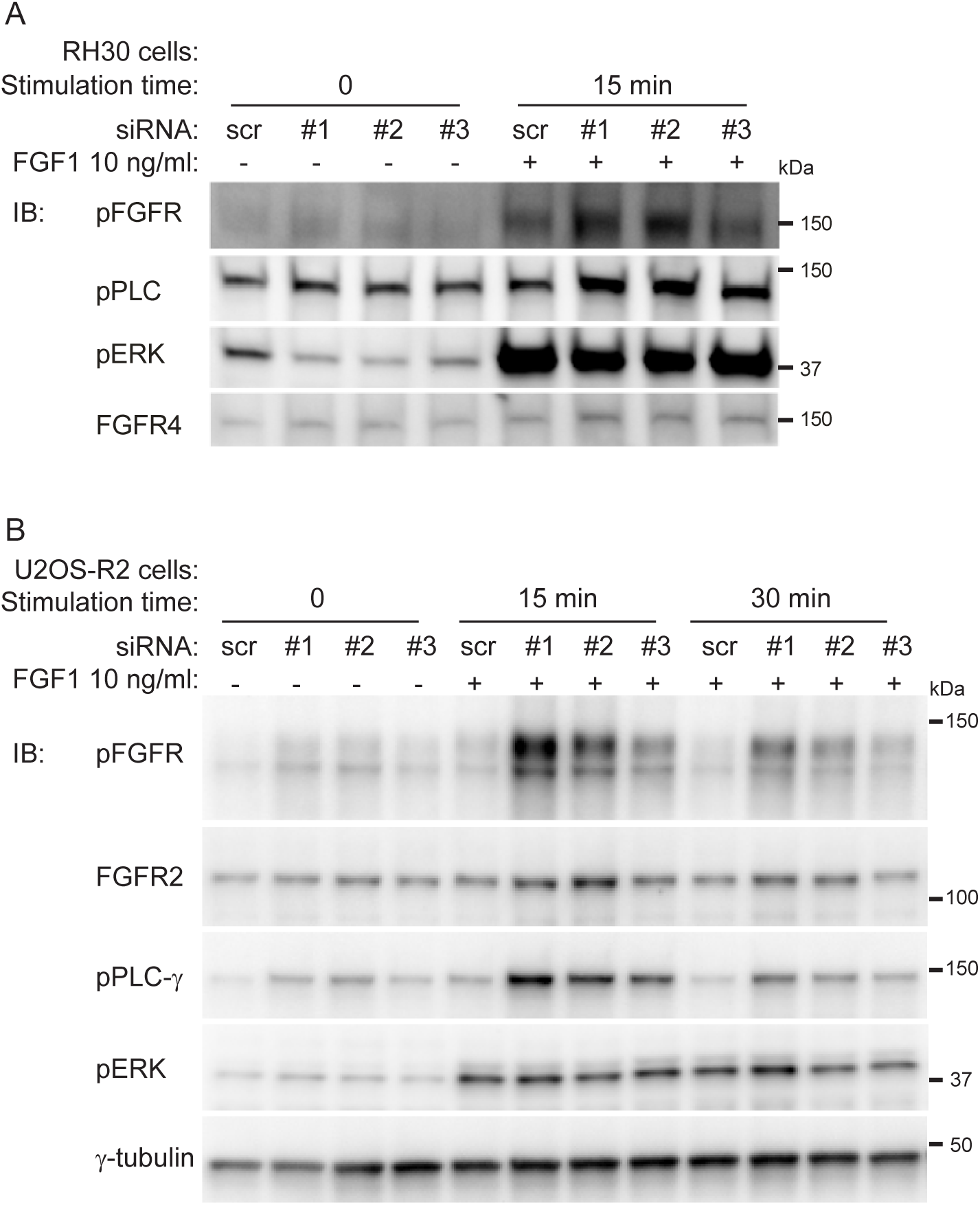
**(A)** RH30 cells were treated with PTPRG siRNAs (#1-#3) or control siRNA (scr) for 72 hours. The cells were then serum-starved for 2 hours before stimulation with FGF1 in the presence heparin for 15 minutes. Next, the cells were lysed and the lysates were subjected to SDS-PAGE followed by western blotting using denoted antibodies. A representative western blot is shown. **(B)** U2OS-R2 cells were treated with PTPRG siRNAs (#1-#3) or control siRNA (scr) for 72 hours. The cells were then serum-starved for 2 hours before stimulation with FGF1 in the presence heparin for 15-30 minutes. Next, the cells were lysed and the lysates were subjected to SDS-PAGE followed by western blotting using denoted antibodies. A representative western blot is shown.

